# Titin cleavage is a driver of cardiomyocyte disengagement and reactive myocardial fibrosis

**DOI:** 10.1101/2025.04.22.645683

**Authors:** Miguel A. López-Unzu, Maria Rosaria Pricolo, Roberto Silva-Rojas, Alejandro Clemente-Manteca, David Sánchez-Ortiz, Natalia Vicente, Diana Velázquez-Carreras, Cristina Morales, Manuel Gavilán-Herrera, Laura Sen-Martín, Ángel Fernández-Trasancos, Verónica Labrador-Cantarero, Lucia Sánchez-García, Carlos Relaño-Rupérez, Fátima Sánchez-Cabo, Laura Lalaguna, Joan Isern, Pura Muñoz-Cánoves, Guadalupe Sabio, Enrique Lara-Pezzi, Elías Herrero-Galán, Jorge Alegre-Cebollada

## Abstract

Myocardial remodeling including cardiomyocyte-death-independent, reactive fibrosis and disconnection of cardiomyocytes is at the basis of prevalent cardiac conditions converging into arrhythmias and heart failure. However, the molecular mechanisms behind these pathogenic responses remain incompletely understood limiting therapeutic opportunities. Here, we find that a molecular event common to unrelated heart diseases, namely the cleavage of the sarcomeric protein titin, is enough to trigger fast myocardial remodeling. Using an engineered system based on the expression of tobacco etch virus protease (TEVp) in mice, we show that 30% mosaic cardiac titin cleavage leads to global cardiomyocyte disengagement, activation of cardiac fibroblasts and interstitial collagen deposition. These effects are concurrent, involve ERK1/2 signaling, and are expected to contribute, at least, to myocardial remodeling in chemotherapy-induced cardiotoxicity and ischemia damage.

---

Excessive extracellular accumulation of collagen (i.e. fibrosis) and alteration of cardiomyocyte-cardiomyocyte junctions are pathological myocardial remodeling processes that can predict adverse outcomes in a variety of heart diseases of different etiology (*1–3*). While the mechanisms driving reparative fibrosis induced by cardiomyocyte death are relatively well known, much less is understood about reactive, interstitial fibrosis occurring in the absence of cardiomyocyte loss (*1, 4*). For instance, how reactive fibrotic responses are mounted during anthracycline-induced cardiotoxicity, or in the surviving, remote myocardium in ischemic accidents, remains elusive (*5, 6*). It is also unclear whether mechanistic hierarchies between fibrosis and cardiomyocyte disengagement exist (*7, 8*). This general lack of understanding is in line with limited efficacy of therapies aimed to prevent detrimental myocardial remodeling (*1, 9*).

Protease activation in cardiomyocytes has been recently proposed to contribute to the development of reactive fibrosis and subsequent decline of cardiac function in chemotherapy-related cardiotoxicity (*10*), myocardial infarction (*11*), the metabolic syndrome (*12*) and left-ventricular pressure-overload stress (*13*). However, the mechanisms linking proteolytic activity and pathogenic myocardial remodeling remain obscure. Here, we hypothesized that cleavage of titin, a fundamental giant polypeptide that scaffolds contractile sarcomeres (*14, 15*) and whose malfunctioning causes hereditary fibrotic heart disease (*16–19*), could be a trigger of myocardial remodeling. In support of this hypothesis, titin cleavage is observed in several unrelated conditions of the heart that result in myocardial fibrosis (*10, 11, 20, 21*) (**Figure 1A**). Importantly, considering the prominent mechanical role of the protein (*22*), a single cleavage event affecting any of the >30,000 peptide bonds in titin could result in similar pathophysiological consequences derived from the unloading of the protein (*23, 24*).

**Figure 1.**
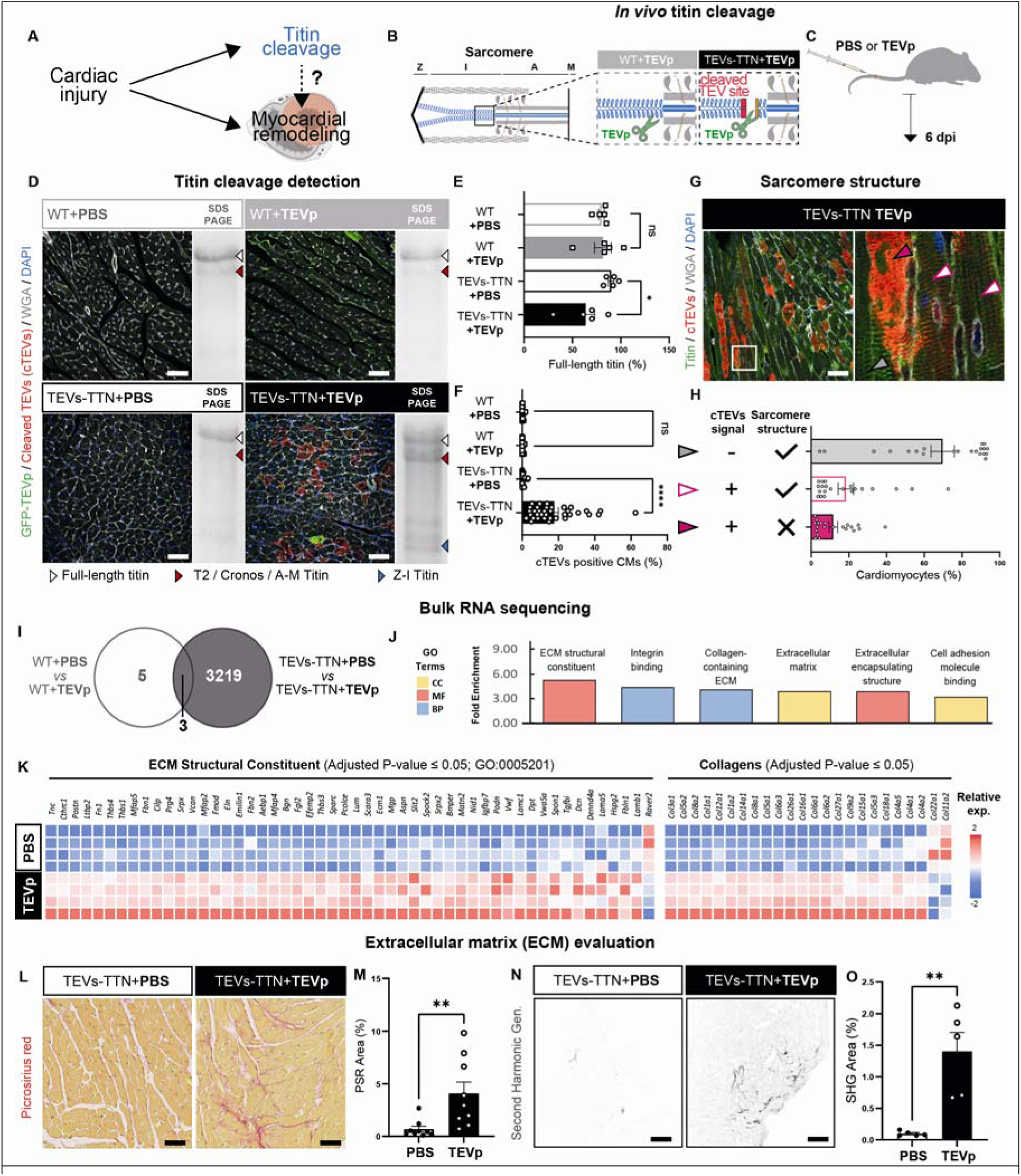
Reactive fibrosis in the myocardium upon titin cleavage. (**A**) Many injuries to the heart result in titin cleavage and myocardial remodeling. Here, we test if titin cleavage is enough to trigger myocardial remodeling. (**B**) Experimental design. Animals from WT and TEVs-TTN backgrounds were injected with PBS (control) and AAVMYO to transfect GFP-TEVp. (**C**) After 6 dpi animals were sacrificed, hearts were sampled and their ventricular myocardia were analyzed. TEVp activity only occurs in TEVs-TTN background. (**D**) Location of GFP-TEVp and cTEVs in cross-sectional cryosections and corresponding titin cleavage detection by SDS-PAGE. Identity of titin bands is indicated at the bottom. (**E**) Quantification of the proportion of full-length titin (N = 5 per group). (**F**) Quantification of proportion of cTEVs-positive cardiomyocytes (N = 8 – 9 per group). (G) Immunofluorescence to test sarcomeric integrity in untargeted (cTEVs negative) and targeted (cTEVs positive) cardiomyocytes in cryosections showing longitudinally oriented cardiomyocytes using fluorescent HaloTag ligand to mark titin (N = 2 animals). Three different morphotypes of cardiomyocytes are found: cTEVs-negative with preserved sarcomeric structure (gray arrowhead), and cTEVs-positive cardiomyocytes with mostly preserved (white arrowhead) or disrupted (pink arrowhead, 11.8%) sarcomeres. (H) Quantification of the three cardiomyocyte morphotypes. (I) Venn diagram showing myocardial DEGs by RNAseq in both WT and TEVs-TTN mice upon infection with TEVp-expressing AAVMYO. (N = 4 per group). (J) Pathway enrichment analysis of the DEGs in the TEVs-TTN group. (K) Relative expression of DEGs (p-value < 0.05) in TEVs-TTN belonging to the GO term “ECM Structural components” (left) and “Collagens” (right). (L) Representative Picrosirius-red stained sections to quantify total collagen in the myocardium. (M) Quantification of the area occupied by collagen (N = 9 per group). (N) Representative images of SHG signal in the myocardium. (O) Quantification of SHG signal. (N = 5 per group). All scale bars = 50 μm. The results are expressed as mean ± standard error of the mean (SEM). p-value <0.05 (*), <0.01 (**),<0.0001 (****). DEGs: differentially expressed genes; dpi: days post-infection; WGA: wheat germ agglutinin. Schemes adapted from www.biorender.com.

## Cardiac titin cleavage triggers rapid interstitial myocardial fibrosis

To test our hypothesis, we examined the consequences of inducing a single, controlled proteolytic cut in cardiac titin. With this aim, we expressed tobacco etch virus protease (TEVp), which is not naturally present in mammals and shows well-defined sequence-dependent activity (*25*), in adult, knock-in TEVs-TTN mice containing the TEV recognition site (TEVs) in the I-band of titin (**Figure 1B,C**) (*26*). Consistent with *in vitro* and *ex vivo* work (*26–28*) using SDS-PAGE we find 30% reduction of full-length cardiac titin and concomitant increase of Z-I and A-M fragments 6 days post infection (dpi) with 1.6-2.6 x 10^12^ GFP-tagged-TEVp-expressing AAVMYO particles (*29*) (**Figure 1D,E, Table S1, Supplementary Data S1**). Titin molecules remain intact in control AAVMYO-injected wild-type (WT) counterparts and in TEVs-TTN mice injected with PBS (**Figure 1D,E**). Notably, the pattern and extent of cardiac titin cleavage we achieve by transducing TEVp in TEVs-TTN mice are similar to the ones resulting from MMP-2 protease activation after anthracycline-induced cardiotoxicity (*10*) and myocardial infarction (*11*).

Staining transverse myocardial sections with anti-cleaved TEV site (cTEVs) antibodies (*23*) reveals a mosaic pattern with an average 18% targeted cardiomyocytes in AAVMYO-injected TEVs-TTN mice (red cells, **Figure 1D,F**), indicating higher sensitivity of anti cTEVs antibodies than anti-GFP counterparts (**Figures 1D, S1A**). Remarkably, almost half of the ∼30% cTEVs-positive cardiomyocytes in complementary longitudinal sections show extensive disruption of sarcomere structure (**Figure 1G,H**), which, however, does not result in increased apoptosis or necrosis (**Figure S1B,C**).

Having validated the TEVp-based system to cleave cardiac titin, we analyzed global transcriptional effects induced by titin cleavage (**Figure 1I-K**). Comparative gene expression analyses reveal that only 8 genes are differentially expressed following transduction of TEVp in WT animals at 6 dpi, indicating that the bare expression of the protease has minor, if any, non-specific effects in the absence of knocked-in TEV sites (**Figure 1I**). The high specificity of the system is further captured by principal component analysis (PCA), which also indicates some level of basal transcriptional differences between WT and TEVs-TTN animals (**Figure S1D**). In sharp contrast, expression of TEVp in TEVs-TTN mice results in major alterations of transcription including 3,222 differentially expressed genes (DEGs, **Figures 1I, S1B**). Differential enrichment analysis in the TEVs-TTN group shows that many of the DEGs belong to categories related with the extracellular matrix (ECM) and cell adhesion (**Figure 1J**). Indeed, the vast majority of DEGs in the GO-term “ECM structural constituents” are clearly upregulated in AAVMYO-injected TEVs-TTN samples (**Figure 1K**). Similar results are obtained with DEGs codifying for collagen proteins (**Figure 1K**). In combination, these results strongly suggest that titin cleavage causes the myocardium to mount a fast reactive fibrotic response. Indeed, at 6 dpi we find increased collagen-marking picrosirius red staining in the myocardial interstitium of AAVMYO-injected TEVs-TTN animals (**Figures 1L,M, S1E**), which also shows higher levels of second-harmonic generation (SHG) signal indicative of well-organized collagen (*30*) (**Figure 1N,O**). This remarkably rapid fibrotic response occurs even as soon as 3 dpi at the highest doses of AAVMYO particles tested (3.8 x 10^12^ viral particles, **Figure S2A, Supplementary Data S1**), and when GFP-TEVp is expressed under a cardiac troponin T (cTnT), cardiomyocyte-specific promoter (**Figure S2B-E**) instead of the constitutive CMV counterpart used in the rest of our experiments.

## Cardiac fibroblasts proliferate and get activated following titin cleavage

Since fibroblasts are the main cell type involved in ECM remodeling (*1*), we examined by immunofluorescence how they react to titin cleavage (**Figure 2A-J**). Using vimentin as a fibroblast marker and alpha smooth muscle actin (αSMA) as an indicator of their activated state (*31*), we observe higher levels of activated fibroblasts in the myocardium of AAVMYO-injected TEVs-TTN animals (**Figures 2A-D, S3A-E**). In the subepicardial zone, these vimentin and αSMA double-positive cells appear typically rounded and isolated (**Figure 2B**), whereas in the subendocardium and innermost layers of the myocardium they develop more stellate cytomorphology (**Figure 2D**). In addition, we find increased proliferation of fibroblasts in AAVMYO-injected TEVs-TTN myocardium according to the higher expression of Ki-67, a marker of cell cycle progression (*32*) (**Figure 2F-J, S3F-J**).

**Figure 2.**
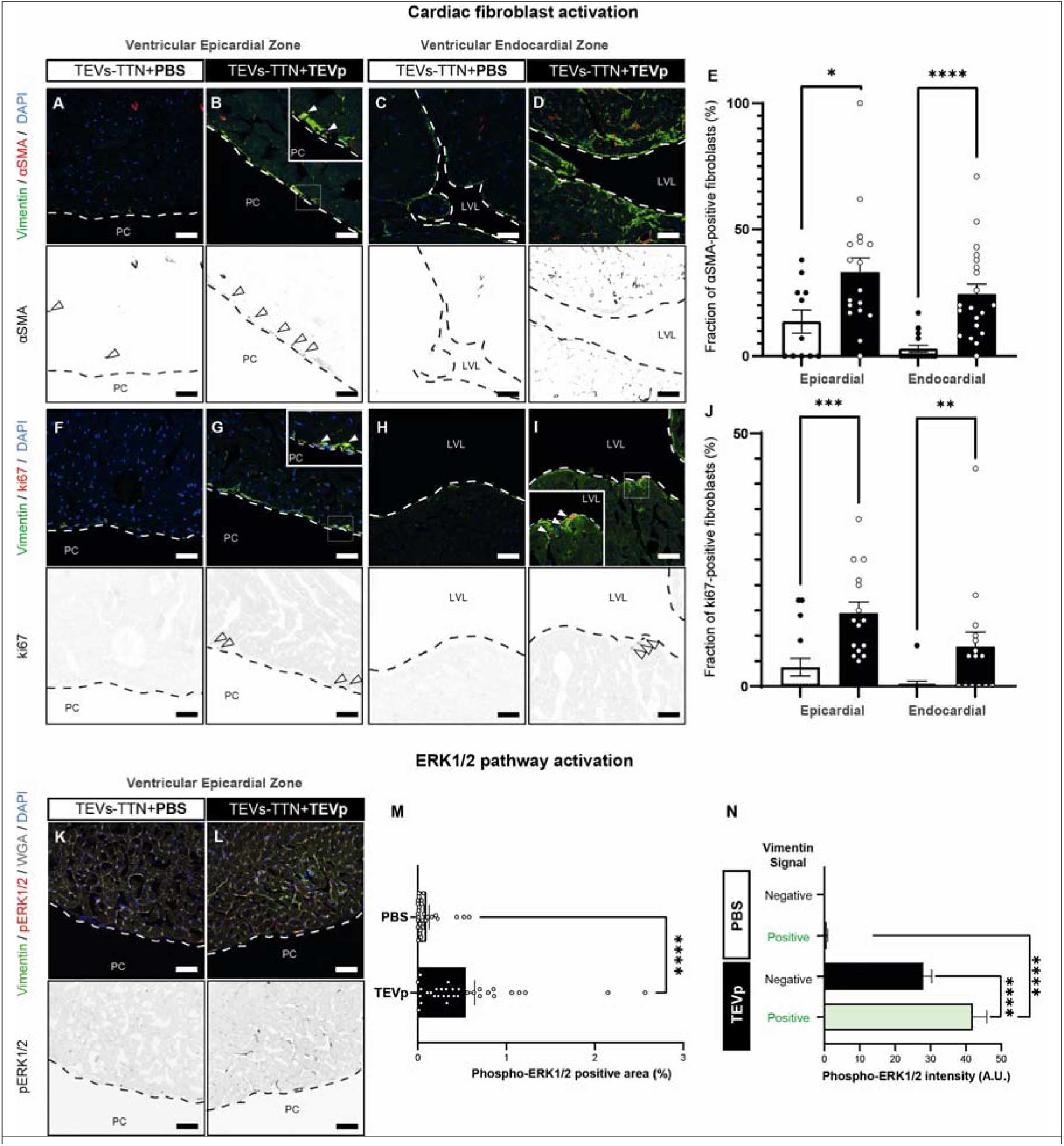
Activation of cardiac fibroblasts upon titin cleavage. (**A-E**) Immunolocalization and quantification of vimentin-positive fibroblasts expressing αSMA in the indicated regions of the myocardium (N = 5 per group). (**F-J**) Immunolocalization and quantification of vimentin-positive fibroblasts expressing Ki67. (**K,M**) Immunodetection and quantification of phospho-ERK1/2 (N = 11 per group). (**N**) Quantification of phospho-ERK1/2 intensity according to vimentin signal. All scale bars = 50 µm. Results are expressed as mean ± standard error of the mean (SEM). p-value < 0.05 (*), <0.01(**), <0.001 (***), <0.0001 (****), non-significant (N.S.). αSMA: alpha-smooth muscle actin; PC: pericardial cavity; LVL: left-ventricle lumen.

In combination, our results so far indicate that titin cleavage results in a fast fibrotic response in the myocardium that occurs in the absence of cell death, and is characterized by cardiac fibroblast proliferation and activation and increased interstitial collagen deposition. Surprisingly, titin-cleavage-induced fibrosis occurs in the absence of increased SMAD2/3 complex phosphorylation, indicating lack of activation of the canonical TGF-β pathway, arguably the main converging route for myocardial fibrosis and cardiac fibroblast activation (*1*) (**Figure S4A**). Indeed, we detect no expression of the canonical transcriptional program associated with TGF-β by RNAseq in AAVMYO-injected TEVs-TTN samples (**Figure S4B**). Similarly, in these samples there is no obvious stimulation of p38, a stress kinase that also has profibrotic functions in the myocardium (**Figure S4C**). In contrast, we find increased phosphorylation of ERK1/2, another stress kinase that has been linked to myocardial fibrosis, in the interstitial zone between cardiomyocytes in the outermost regions of the ventricles of titin-cleaved-containing myocardium (**Figure 2K-M**). Accordingly, RNA-seq data captures activation of target genes downstream of ERK1/2 (**Figure S4D**). Of note, we find that the phospho-ERK1/2 signal is higher in areas that are also vimentin positive, indicating activation of ERK1/2 in fibroblasts (**Figure 2N**).

## Titin cleavage directly compromises cardiomyocyte-cardiomyocyte junctions

Our RNAseq data suggest alteration of cardiomyocyte adhesion upon titin cleavage (**Figure 1J**). To validate this observation, we stained histological sections using antibodies against connexin 43, a main component of gap junctions in the intercalated discs of cardiomyocytes that ensures electrical coupling and that is highly sensitive to perturbed cardiomyocyte-cardiomyocyte adhesion (*2*). Results show extensive, global reduction of myocardial connexin 43 levels in the intercalated discs of AAVMYO-injected TEVs-TTN animals (**Figure 3A,B**), indicating major loss of cardiomyocyte engagement upon titin cleavage. Intriguingly, the loss of connexin 43 occurs both in cTEVs-positive and negative areas, although it is more intense in cTEVs-negative regions (**Figure 3B**).

**Figure 3.**
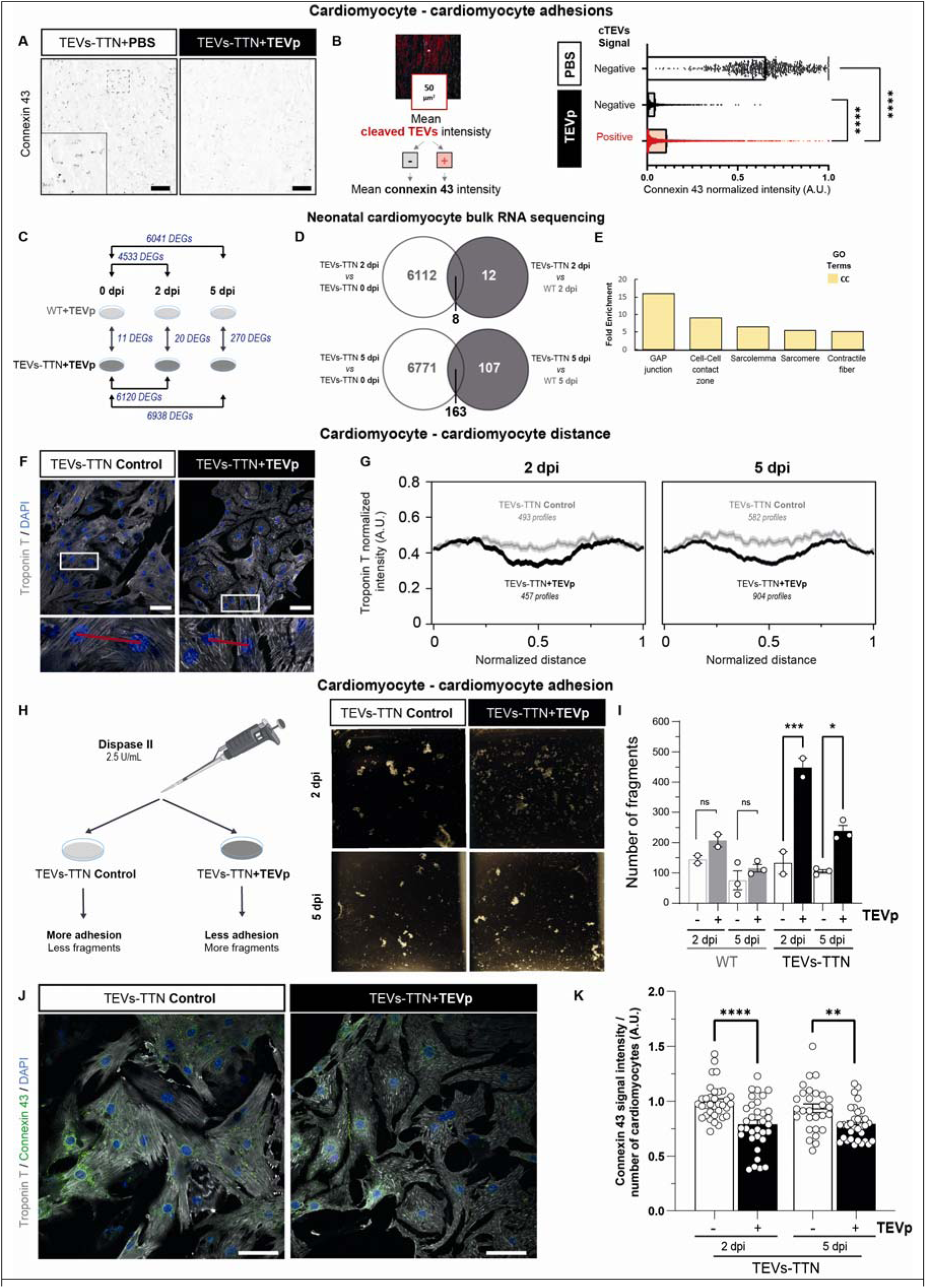
Disruption of cardiomyocyte-cardiomyocyte adhesion following titin cleavage. (**A**) Representative images showing connexin 43 immunofluorescence. **(B)** Quantification of connexin 43 signal intensity in cTEVs negative and positive areas in control and AAVMYO injected samples. (C) Summary of DEGs obtained by RNAseq using cultured neonatal cardiomyocytes (N = 4 per group). (D) Venn diagrams showing overlapping DEGs obtained in the indicated comparisons. (E) GO analysis of the 163 DEGs found in both TEVs-TTN 5 dpi vs TEVs-TTN 0 dpi vs and TEVs-TTN 5 dpi vs WT 5 dpi. (F) Evaluation of cardiomyocyte-cardiomyocyte contacts from the intensity of troponin T immunofluorescence signal between nuclei (red lines) (G) Average troponin T fluorescent signal between nuclei (N = 3 independent experiments). (H) Dissociation assay reporting on cardiomyocyte–cardiomyocyte adhesion and representative images of resulting fragments. (I) Number of fragments in the dissociation assay (N = 2-3 independent experiments per condition). (J) Immunostaining of gap junction protein connexin 43 in neonatal cardiomyocytes. (K) Quantification of connexin 43 intensity, which was normalized by the number of cardiomyocytes in the image. A minimum of N = 30 cells per condition were quantified in a total of 3 independent experiments. Signal intensities were normalized considering the maximum fluorescence measured in each of the three experiments. Scale bars: 100 μm (A) or 50 μm (F,J). Results are expressed as mean ± standard error of the mean (SEM). p-value < 0.05 (*), <0.01(**), <0.001 (***), <0.0001 (****). DEG: differentially expressed gene; GO: gene ontology. Schemes adapted from www.biorender.com.

To dissect whether cardiomyocyte disengagement upon titin cleavage is independent of fibrosis, we expressed TEVp in isolated neonatal TEVs-TTN cardiomyocytes using AAV6 vectors (*28*). We did bulk RNAseq of AAV6-infected cardiomyocytes at different times post infection, including also WT cells as controls (**Figure 3C-E**). In non-infected cells (0 dpi), only 11 genes are differentially expressed between WT and TEVs-TTN samples, indicating that basal phenotypical differences between genotypes are small in neonatal cardiomyocytes. At 2 and 5 dpi, we find that the transcriptomes of both WT and TEVs-TTN cardiomyocytes change considerably (**Figure 3C**), an effect most probably contributed by aging of cultured cardiomyocytes. We enriched transcriptional changes associated specifically to titin cleavage by selecting common DEGs in infected TEVs-TTN cardiomyocytes when compared both to non-infected counterparts and with infected WT samples at the same dpi (**Figure 3D**). This analysis identifies 8 and 163 DEGs at 2 and 5 dpi, respectively, which are time points when the majority of titin molecules have been cleaved (*28*). Remarkably, enrichment analysis identifies that these transcriptional changes primarily affect genes involved in sarcomere function and in cardiomyocyte-cardiomyocyte junctions (**Figure 3E**).

We validated perturbed cardiomyocyte-cardiomyocyte adhesion in the *in vitro* system by several approaches. First, we analyzed inter-cardiomyocyte space by monitoring the immunofluorescence signal for intracellular troponin T in lines connecting cardiomyocyte nuclei (example connecting lines shown in red, **Figure 3F**). Averaging >450 intensity profiles per condition, we find reduced troponin T signal in the mid region between nuclei in AAV6-infected TEVs-TTN samples as early as 2 dpi, a result that is compatible with titin cleavage rapidly inducing less intimate contact between cardiomyocytes (**Figure 3G**). In agreement with this observation, AAV6-infected TEVs-TTN display reduced intercellular adhesive forces, as determined by increased number of fragments detected in dissociation assays where confluent cell monolayers are exposed to mechanical stress (**Figure 3H,I**) (*7*). Further confirming cardiomyocyte-cardiomyocyte disconnection, we observe reduced connexin 43 signal intensity by immunofluorescence in AAV6-infected TEVs-TTN cardiomyocytes (**Figure 3J,K**).

## Titin cleavage boosts interaction with the ECM via expression of integrin **α**5-**β**1

Our *in vitro* experiments demonstrate that titin cleavage alters cardiomyocyte connectivity in a manner that is independent of fibrosis. Remarkably though, the *in vivo* results indicate that even when titin cleavage is induced in no more than 20-30% cardiomyocytes, we observe global cardiomyocyte-cardiomyocyte disengagement that also involves cells with intact titin (**Figure 3A,B**). We asked ourselves whether loss of cell connectivity is equivalent in cardiomyocytes containing cleaved titin and in non-targeted counterparts. To address this question, we compared mRNA expression in single-nuclei from TEVs-TTN ventricular myocardia infected or not with GFP-TEVp-expressing AAVMYO. Using UMAP representation to reduce dimensionality, we observe that the expression profile of nuclei from cardiomyocytes differs considerably between both experimental groups (**Figures 4A, S5**). Indeed, we identify one major cluster of cardiomyocyte nuclei (*CM1*) that is mostly found in control conditions and nearly absent in AAVMYO-TEVp injected TEVs-TTN samples (**Figure 4B**). In contrast, three additional cardiomyocyte clusters (*CM2-CM4*) are almost exclusive of AAVMYO-injected TEVs-TTN samples (**Figure 4B**). Hence, we reasoned that these three clusters probably derive from *CM1*. Indeed, RNA velocity analysis (*33*) captures two main transitions originating from CM1 and ending, respectively, in *CM2* and *CM4* (**Figure 4C**). *CM3* appears as an intermediate cluster in the trajectory from *CM1* to *CM4* (**Figure 4C**).

**Figure 4.**
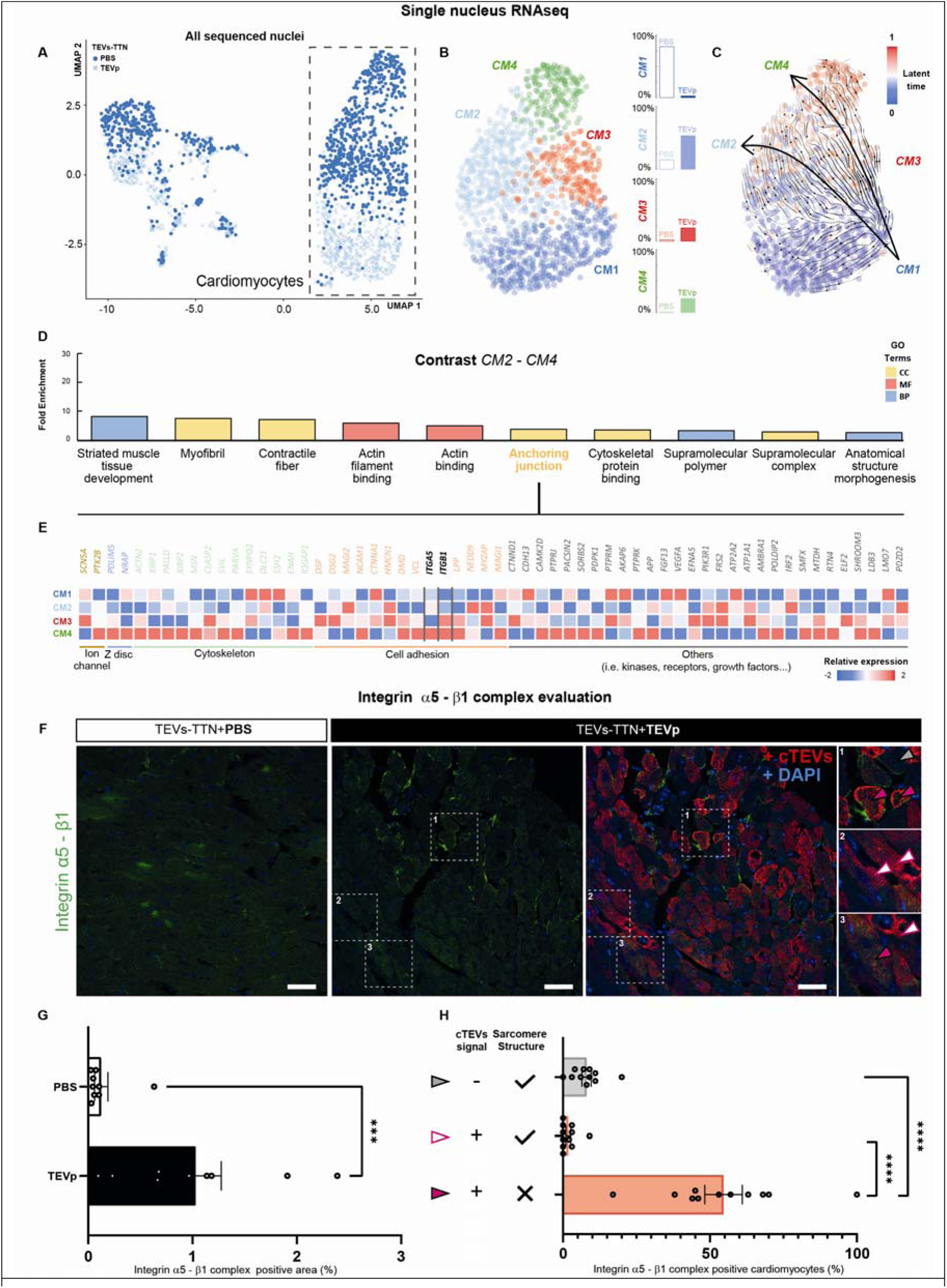
Single-nuclei RNAseq analysis of mosaic titin cleavage. (**A**) UMAP representation of single nucleus RNA sequencing of myocardial samples (N = 4 per group; PBS, light blue crosses; TEVp, blue circles). (**B**) Clustering of cardiomyocyte populations, annotated as *CM1* (dark blue), *CM2* (cyan), *CM3* (red) and *CM4* (green). (C) RNA velocity analysis returns two possible trajectories between CM clusters: one from CM1 to CM2 and other from CM1 to CM4 via CM3. (D) Pathway enrichment analysis of DEGs in the contrast between CM2 and CM4. The top dysregulated terms, selected by decreasing FDR and ordered in the top 10 by their fold-enrichment, are related to the cytoskeleton and myofibril. The analysis also returns the GO term “anchoring junction” (GO: 0070161). (E) DEGs within this term are associated with different structures of these adhesions (ion channel, cytoskeleton or Z-discs) or with cellular adhesion itself. Among these, integrins ITGA5 and ITGB1 stand out as they are the only elements connecting the extracellular matrix and the cardiomyocyte membrane that are highly upregulated in CM4. (F) Immunolocalization of integrin α5–β1 complex and cTEVs. See panel H for explanation of symbols. (G) Quantification of integrin α5 – β1 complex signal (N = 2 – 3 per group). (H) Fraction of integrin α5–β1-positive cardiomyocytes according to cTEVs positivity and structural integrity of sarcomeres (N = 2 – 3 per group). cTEVs- and integrin-α5–β1-negative cells are identified from WGA staining (not shown). All scale bars = 50 μm. The results are expressed as mean ± standard error of the mean (SEM). p-value <0.001 (***), <0.0001 (****). UMAP: uniform manifold approximation and projection; DEG: differentially expressed gene;

Considering the mosaic nature of TEVp activity in AAVMYO-injected TEVs-TTN myocardium, we reasoned that one of the two final clusters of cardiomyocytes (i.e., *CM2* or *CM4*) could be enriched in non-targeted cells while the other could be contributed by cardiomyocytes containing cleaved titin. We examined this scenario by doing pathway enrichment analysis considering DEGs between clusters *CM2* and *CM4*. The main differences captured by this analysis are found in terms related to muscle contraction and development, the cytoskeleton and cell adhesion (**Figure 4D**). Among the latter, there is clear upregulation of integrins α5 and β1 in cluster *CM4* compared with the remaining clusters (**Figure 4E**). Accordingly, we find increased levels of integrin α5-β1complex in AAVMYO-injected TEVs-TTN samples (**Figure 4F,G**). Notably, over 50% of cardiomyocytes that are cTEVs-positive and show disrupted sarcomeres in AAVMYO-injected TEVs-TTN myocardium are also positive for expression of integrin α5-β1 complex (**Figure 4F,H**), strongly suggesting that cardiomyocytes containing cleaved titin and extensively affected sarcomeres are main contributors to the *CM4* cluster. This situation would imply that *CM3*, a cluster transitioning towards *CM4*, most probably is enriched in cardiomyocytes containing cleaved titin but preserved sarcomeres, which would leave *CM2* as a population of cardiomyocytes with intact titin molecules in line with the diverging trajectories between *CM2* and *CM4* detected by RNA velocity analysis (**Figure 4C**).

Integrin α5-β1, which strongly binds fibronectin in the ECM (*34*), is expressed prevalently in fetal and neonatal cardiomyocytes with immature sarcomeres (*35*), but also in situations of stress including ischemia (*36*). Overexpression of integrin α5-β1 could shift transmission of myocardial mechanical stress from cardiomyocyte-cardiomyocyte junctions to cardiomyocyte-ECM adhesion sites contributing to preservation of myocardial morphology in the absence of fully functional cell adhesions (*37*). Our single nuclei RNAseq and histological data indicate that this response in TEVs-TTN myocardium is cell-autonomous and restricted to cardiomyocytes containing cleaved titin.

## Cardiomyocyte disengagement and fibrosis responses upon titin cleavage are concurrent

Next, we aimed at studying the kinetics of the main remodeling events that are triggered by titin cleavage. We set up an independent cohort of animals injected with a standard dose of GFP-TEVp-expressing AAVMYO that were analyzed at 3 dpi (**Table S1**). At this early time point, full-length titin is reduced by only 18% (**Figure S6A**) and cTEVs staining above background levels is observed in just 4% cardiomyocytes (**Figure S6B**). Despite this limited titin cleavage, AAVMYO-injected animals show already increased interstitial ERK1/2 signaling (**Figure S6C**) and a quite remarkable fraction of proliferative fibroblasts (**Figure S6D**). These events are not accompanied by overt fibrosis yet (**Figure S6E**); however, we find a tendency for increased activation of fibroblasts in the targeted myocardium (**Figure S6F**). Similarly, we observe alterations of connexin 43 levels in cardiomyocytes, which are transiently elevated at this time point in AAVMYO-injected TEVs-TTN animals (**Figure S6G**), as well as increased levels of integrin α5-β1 (**Figure S6H**). These results demonstrate that the myocardial remodeling processes triggered by low levels of titin cleavage start very rapidly.

## Discussion

Here, we demonstrate that titin cleavage in a limited fraction of cardiomyocytes is sufficient to trigger one of the fastest, if not the fastest, myocardial reactive fibrotic responses identified so far (*38*). Remarkably, full loss of titin has been observed to cause severe heart disease, but no myocardial fibrosis (*39*), suggesting that the profibrotic response described here is not secondary to sarcomere disassembly (*23*) but results instead from toxicity of unloaded titin molecules. At the mechanistic level, we have found increased signaling through ERK1/2 correlating with higher proliferation and activation of cardiac fibroblasts. Surprisingly, titin-cleavage-induced fibrosis does not result in noticeable activation of p38 or the canonical TGF-β pathways, which are typical profibrotic signals in the myocardium that are induced for instance by the renin-angiotensin-aldosterone system, during inflammatory responses or when cardiomyocyte desmosomes are compromised (*1, 7*).

Regarding the alteration of cardiomyocyte-cardiomyocyte adhesion we have observed upon titin cleavage, we speculate that the loss of force transmission at transitional junctions following unloading of titin (*40*) could weaken catch-bond-based cadherin adhesions (*41*) and subsequently impair gap junctions (*42*). Interestingly, we detect increased ERK1/2 activation even when loss of connexin 43 has not been completed, suggesting that both reactive fibrosis and disengagement of cardiomyocytes are not hierarchical but instead occur concurrently. In support of this scenario, loss of cardiomyocyte adhesion in our *in vitro* experiments occurs in the absence of fibrosis, and cardiac-restricted connexin 43 knock-out mice do not develop myocardial fibrosis (*43*).

From a translational perspective, our results imply that equivalent remodeling processes to the ones observed in TEVp-expressing TEVs-TTN myocardium should also be present in conditions where titin cleavage is induced by endogenous protease activation, including ischemic disease (*11, 21*), anthracycline-induced cardiotoxicity (*10*) and atrial fibrillation (*20*). Remarkably, perturbed connexin 43 levels (*44–46*), activation of ERK1/2 (*47–49*) and reactive fibrosis (*3, 5, 49*) have all been found in these conditions, strongly suggesting that titin cleavage plays a previously unappreciated mechanistic role in myocardial remodeling processes underlying different forms of heart disease. Accordingly, inhibition of MMP-2 prevents both cleavage of titin and myocardial fibrosis in experimental models of anthracycline-induced cardiotoxicity (*10*). Furthermore, we speculate that arrhythmic substrates typical of the aforementioned conditions (*50, 51*) may be contributed by the combination of fibrosis and connexin 43 alteration caused by titin cleavage (*1, 52*). Indeed, the dramatic loss of connexin 43 and the fibrosis found in targeted TEVs-TTN myocardium are accompanied by noticeable changes in electrophysiological parameters (**Figure S7**) and high mortality in the absence of major systolic or diastolic cardiac dysfunction (**Figure S8**).

Looking to the future, our results highlight the key role of titin in acquired heart disease, which adds to the well-known contribution of the protein to hereditary cardiomyopathy. Similar to rodent models of disease, coexisting fibrosis and titin degradation can be observed in terminally failing human hearts (*53, 54*), in ischemic human myocardium (*55*) and in right-ventricular samples from pulmonary hypertension patients (*56*) (**Figure S9**). We suggest that identification of causative proteases in these and other conditions could provide new windows of therapeutic opportunity based on protection of titin integrity. It is also possible that similar titin-based pathomechanisms could contribute to diseases caused by variants in the titin gene. For instance, dilated cardiomyopathy, a familial condition that usually presents with myocardial fibrosis (*57*), is most commonly caused by truncated titins that are prone to recoiling towards the Z-disk of sarcomeres (*58, 59*) and that can therefore trigger profibrotic mechanosignaling similar to cleaved counterparts (*23*). Indeed, truncating titin variants have been specifically associated with the development of primary myocardial fibrosis in individuals with sudden cardiac death

(*16*). Similarly, missense variants that render titin more sensitive to proteolysis cause arrhythmogenic cardiomyopathy, a fibrotic myocardial disease that is a frequent trigger of fatal arrhythmias (*19*). We propose that consideration of downstream events to titin unloading will help overcoming the challenges associated with designing effective treatments for pathogenic myocardial remodeling (*60*). More generally, our work demonstrates that alteration of sarcomere function in otherwise viable cardiomyocytes can rapidly affect the ECM via TGF-β-independent activation of non-cardiomyocyte cells, a so far unnoticed cardiac fibrosis route.

## Acknowledgments

CNIC is supported by the Instituto de Salud Carlos III (ISCIII), the Ministerio de Ciencia, Innovación y Universidades (MCIU, MICIU/AEI/10.13039/501100011033) and the Pro CNIC Foundation, and is a Severo Ochoa Center of Excellence (grant CEX2020-001041-S funded by MCIU). Single-nucleus RNAseq experiments were performed in the Genomics Unit of the CNIC. Light microscopy was conducted at the CNIC Microscopy & Dynamic Imaging Unit. Biomedical Imaging was conducted at the Advanced Imaging Unit of the CNIC using ReDIB ICTS infrastructure TRIMA@CNIC, MCIN with the support of ISCIII (PT20/00044), co-financed by the European Union through the European Regional Development Fund (ERDF, “A way of doing Europe”). We acknowledge the personnel from CNIC animal housing (especially Bahia El Maimouni and Tamara Anguita), viral vectors, genomics, flow cytometry, bioinformatics and histology facilities. We acknowledge the feedback from many colleagues at CNIC, especially Sara Martínez, Juan Miguel Redondo, Juan Bernal, Andrés Hidalgo, Miguel Ángel del Pozo, José Luis de la Pompa, Hesham Sadek, Valentín Fuster, Florian Weinberger and Miguel Torres. We thank Mateo Sánchez for input regarding TEVp. We acknowledge the feedback from Benjamin Prosser, Jose Maria Pérez-Pomares, Adrian Villalba Ruiz and Javier Díez. We thank all members of the Molecular Mechanics of the Cardiovascular System team for their support and input. We thank Alba Pobes-Lagartos for excellent technical assistance.

## Author contributions

Conceptualization: MALU, MRP, RSR, MGH, AFT, EHG, JAC

Methodology: ACM, DSO, NV, DVC, CM, LSM, VLC, LSG, CRR, LL, JI, GS

Investigation: MALU, MRP, RSR, EHG, JAC

Funding acquisition: MALU, RSR, MGH, GS, JAC

Supervision: FSC, PMC, ELP, JAC

Writing – original draft: MALU, MRP, JAC

Writing – review & editing: RSR, ACM, DSO, NV, DVC, CM, MGH, AFT, VLC, LSG, CRR, FSC, LL, JI, PMC, GS, ELP, EHG

## Data and materials availability

The main data supporting the results in this study are available within the paper and its Supplementary Information. Non-commercial materials are available from the corresponding authors. Transcriptomic data from bulk RNA-seq and single nucleus RNA-seq referenced GRCm38.99 transcriptome are available from https://www.ebi.ac.uk/biostudies/ (E-MTAB-14985 and E-MTAB-14993 accession codes for bulk RNAseq in neonatal mouse cardiomyocytes and hearts, respetively, and E-MTAB-14976 for single nucleus RNAseq).

## Supplementary Materials

Materials and Methods

Supplementary Text S1 Figs. S1 to S9

Tables S1 to S2

Data S1 to S2

