## Supplemental Information for "Titin cleavage is a driver of cardiomyocyte disengagement and reactive myocardial fibrosis"

**The PDF file includes:**

Materials and Methods

Supplementary Text S1

Figs. S1 to S9

Tables S1 to S2

**Other Supplementary Materials for this manuscript include the following:**

Data S1 to S2

Materials and Methods

Animal experimentation

Mice were housed in ventilated cages with free access to food and water and 12h day/night cycles. Heterozygous TEVs-TTN animals were crossed to obtain WT and homozygous animals. TEVs-TTN animals were described before (*26*), and 5´-CGTGGTGGCTTATCTTCTAGC-3´ and 5´-CTGTTGGTTCATGCATCTCC-3´ primers were used for genotyping. Only male mice were used for the experiments. Animals were handled in accordance with European and Spanish guidelines for animal welfare and with the recommendations in the Guide for the Care and Use of Laboratory Animals of the National Institutes of Health. The protocol was approved by the Ethics Committee of Animal Experiments of our institutional committees (project numbers PROEX 042/18 and PROEX 107.8/23).

Adeno associated virus (AAV) production

Among available TEV-protease (TEVp) constructs that are expressed under the CMV promoter (*28*), we have used GFP-uTEV3, which contains an improved version of TEVp obtained by directed evolution (*61*). To drive expression of GFP-uTEV3 under a cardiac troponin T promoter, the original vector was modified by restriction/ligation (**Supplementary Text S1**). The production of adeno-associated virus serotype 6 (AAV6) particles involved a dual transfection process of HEK293T-derived cell line (AAVPro®293T; Takara Bio), utilizing a plasmid containing adenovirus helper sequences in conjunction with rep (AAV2) and cap (AAV6) genes (PF0406, PlasmidFactory) along with the plasmid carrying GFP-uTEV3. For AAVMYO particles, we used a modified cap vector based on literature (*29, 62*) and pAdDeltaF6 helper plasmid. Subsequent to cell lysis in 50mM Tris, pH 8, 150 mM NaCl and three freeze/thaw cycles, we treated cells with 150 U/mL Benzonase (Millipore) at 37°C for 30 minutes and clarified by centrifugation (3,000 g, 5 minutes at room temperature). Next, viral particles underwent purification through iodixanol gradient ultracentrifugation in Polypropylene Optiseal tubes (361625, Beckman-Coulter) at 183,000 g, 3 hours at room temperature. Further concentration was achieved using Amicon Ultra-15 tubes (100K MWCO, Millipore). Samples were stored in PBS buffer supplemented with 0.001% pluronic F-68. Quantification of particle abundance employed qPCR targeting the CMV promoter (ACCATGGTGATGCGGTTTTG and ATGGGGTGGAGACTTGGAAATC primers) or eGFP (AGCTGACCCTGAAGTTCATCTG and AAGTCGTGCTGCTTCATGTG primers).

Systemic injections of viral vectors and cardiac tissue processing

WT and homozygous TEVs-TTN mice aged 8 - 26 weeks were injected with AAVMYO viral particles or 100 µl PBS, 0.001% pluronic acid via the tail vein (**Table S1, Supplementary Data S1**). The mice were randomly injected in order to avoid differences arising from the skill of the researcher to inject. Mice were then returned to their breeding cages until sacrifice by CO_2_ inhalation six days post infection (dpi). Sample sizes were estimated according to previous experience from the investigators considering availability of animals and using the least number of animals possible. Upon sacrifice, the heart together with the ascending aorta and pulmonary trunk, were dissected out. The ascending aorta, from the sinotubular junction to the branching of the brachiocephalic artery and atrial myocardium were excised and discarded. Transverse ventricular samples were obtained from which the most apical region was cryopreserved for protein and RNA extraction, the intermediate region was cryopreserved in liquid nitrogen for subsequent extraction of nuclei, and the upper region of both ventricles was fixed in 4% paraformaldehyde (PFA4%) overnight at 4ºC. The fixed PFA4% samples were washed in PBS, treated with 10% and 30% sucrose solutions and embedded in Tissue-Tek® OCT. Compound (4583, Sakura). Eight μm transverse sections of the intermediate region were used for immunofluorescence and quantification of apoptosis, necrosis, interstitial collagen and collagen organization as detailed below.

Quantification of titin cleavage

For homogeneization, a small piece of ventricular myocardium was placed in a TaKaRa® BioMasher Standard tube (9791A, Takara) with 40 µL of protein extraction buffer (50 mM Tris Buffer pH 6.8, 10 mM EGTA, 3% SDS, 50 mM N-ethylmaleimide) per mg of tissue. Next, samples were heated at 60ºC for 10 minutes and the soluble fraction was obtained by centrifugation at 18,000 g for 5 minutes at room temperature. Samples were prepared for SDS-PAGE analysis by diluting protein extracts with 4x Laemmli sample buffer (containing 0.3% (w/v) bromophenol blue and 50% (v/v) glycerol) as previously described (*63*). Samples were loaded onto 2% polyacrylamide / 1.5% agarose gels for electrophoretic separation. Following electrophoresis, gels were stained with Coomassie Brilliant Blue to visualize protein bands. Band intensities were measured using QuantityOne Software (BioRAD).

Neonatal cardiomyocyte isolation and infection with AAV6 particles

Neonatal mouse cardiomyocytes were isolated from 6-8 pups per experimental group. Dissected hearts from one-to-three-day-old mice were minced with a scissor in cold Hanks’ Balanced Salt Solution (HBSS) and dissociated using the Pierce Primary Cardiomyocyte Isolation Kit (Thermo ScientificTMPierceTM, 88281) in 0.21 mL enzyme mix/heart in a 2 mL reaction tube for 20 minutes at 37°C. After centrifugation at 600g for 5 min, the pellet was washed twice with HBSS and resuspended in 0.5 mL cardiomyocyte medium (DMEM for Primary Cell Isolation containing 10% heat-inactivated FBS and 1% penicillin/streptomycin)/heart and plated onto MatTek dishes covered with matrigel solution (Fisher Scientific, 11553620). 24-48h after plating, cardiomyocytes were infected with AAV6-expressing GFP-TEVp at 13,450 multiplicity of infection (*28*). Media were replaced with cardiomyocyte medium 16 h after virus addition.

RNA isolation and bulk sequencing

RNA was isolated from hearts stored at -80ºC in 500 μL of RNAlater™ Stabilization Solution (Invitrogen, Cat. No. AM7020) homogenized with T10 basic ULTRA-TURRAX® (IKA) or from neonatal cardiomyocytes cultured on 24-well plates (Mattek, P24G-1.5-13-F). Total RNA was isolated by using Trizol (ThermoFisher Scientific) followed by chloroform precipitation and a purification step using RNeasy micro-Kit (Qiagen, Cat. No. 74004) following the indications of the provider. RNA quantity was quantified from its absorbance at 260 nm, while purity was assessed from the ratios of absorbance at 260 nm and 280 nm, and between 260 nm and 230 nm. Sample quality was evaluated using RNA Integrity Number (RIN), which was measured using a 6000 Nano chip in a 2100 Bioanalyzer (Agilent Technologies). For next generation sequencing (NGS) experiments, only RNA samples with RIN > 9 were used. Messenger RNA was purified from total RNA using poly-T oligo-attached magnetic beads. After fragmentation, the first strand cDNA was synthesized using random hexamer primers followed by second strand cDNA synthesis. The library was ready after end repair, A-tailing, adapter ligation, size selection, amplification, and purification. The kit for library preparation was the Novogene NGS RNA Library Prep Set (PT042). The library was checked with Qubit 2.0 and real-time PCR for quantification, and a bioanalyzer 2100 for size distribution detection. Quantified libraries were pooled and sequenced on the Illumina platform Novaseq6000, according to effective library concentration and data amount (6 Gb). The sequencing strategy was pair-end 150 bp (PE150). The clustering of the index-coded samples was performed according to the manufacturer’s instructions. After cluster generation, the library preparations were sequenced on the Illumina platform Novaseq6000, and paired-end reads were generated.

For myocardial samples, raw reads in fastq format were first processed through in-house Perl scripts. In this step, clean reads were obtained by removing reads containing adapter or poly-N, as well as low-quality reads. At the same time, Q20, Q30, and GC content of the clean data were calculated. Reference genome and gene model annotation files were directly downloaded from the genome website. Index of the reference genome was built using Hisat2 v2.0.5 (*64*) and paired-end clean reads were aligned to the reference genome using Hisat2 v2.0.5. FeatureCounts v1.5.0-p3 (*65*) was used to count the read numbers mapped to each gene. Next, the fragments per kilobase of transcript per million mapped reads of each gene was calculated based on the length of the gene and the read counts mapped to this gene. Differential expression analysis of two conditions/groups was performed using the DESeq2R package (1.20.0) (*66*). DESeq2 provide statistical routines for determining differential expression in digital gene expression data using a model based on the negative binomial distribution.

For neonatal cardiomyocytes samples, Sequencing reads were pre-processed by means of a pipeline that used FastQC (*67*), to assess read quality, and Cutadapt (*68*) to eliminate adaptor remains and to discard reads that were shorter than 30 bp. Resulting reads were mapped against reference transcriptome GRCm38.99 and quantified with RSEM (*69*), using Bowtie as aligner. Expected expression counts calculated with RSEM were processed with an analysis pipeline that used the Bioconductor package Limma (*70*) for normalization and differential expression testing, taking only into account those genes expressed with at least 1 count per million in at least two samples.

The resulting p-values were adjusted using the Benjamini and Hochberg's approach for controlling the false discovery rate. Genes with an adjusted p-value ≤ 0.05 were assigned as differentially expressed. For enrichment analysis by Gene Ontology (GO) we have used PANTHER 19.0. We used bar plots analysis considering all differentially expressed genes (DEGs) thresholds to represent the top enriched Cellular Component (CC), Biological Process (BP) and Molecular Function (MF) GO-Terms. For enrichment analysis of specific pathways (i.e. TFG beta pathway) we have used Gene set enrichment analysis (GSEA) showing the results using a enrichment plot (*71*).

Immunohistochemistry

Fixed hearts were embedded in optimum cutting temperature (OCT, Tissue-Tek® Compound 4583, Sakura) and sectioned at 8 μm with a cryostat (Leica CM-1950). Transverse sections of the middle portion of each segment were permeabilized with 0.3% Triton X-100 in PBS dilution. After washing with PBS, non-specific binding sites were saturated for 30 min with 10% FBS, 1.5% bovine serum albumin (BSA) in PBS (SB). Primary antibody incubations were performed at 4°C overnight (see **Table S2** for a list of antibodies used in this work). Primary antibodies were diluted 1:100. After incubation, the slides were washed in PBS, incubated for 1.5 h at room temperature in secondary Alexa Fluor-conjugated anti-rabbit IgG, anti-mouse IgG, anti-goat IgG, or anti-chicken IgY (Sigma-Aldrich) antibodies diluted 1:500 in PBS. Sections were also incubated in 1.0 mg/mL wheat germ agglutinin (WGA) labelled with Alexa Fluor (W32466, ThermoFisher Scientific) diluted 1:250 in PBS for labelling cardiomyocyte cell membranes in histological sections. HaloTag Ligands® conjugated with Alexa-Fluor 488 (Promega) at 0.1 μM, were used in TEVs-TTN samples to label the I-band region of titin (*26*). Negative controls did not include primary antibodies. After the secondary antibody incubation, sections were stained with 0.5 μg/mL 4′,6-diamidino-2-phenylindole dihydrochloride (DAPI, Merck) to mark the location of the nuclei. Mowiol 4-88 (Sigma-Aldrich) was used as mounting medium and allowed to cure overnight. Images were taken using a Nikon A1R confocal microscope (Nikon, Japan) with a Plan Apo 40x/1.3 Oil objective.

Immunofluorescence analysis

The area occupied by positive signal, the signal intensity and the number of positive areas were quantified in microphotographs using ImageJ-FIJI (*72*). Additionally, we have used Cellpose (*73*) wrapper plugin (https://github.com/BIOP) for general cellular segmentation, Stardist plugin for nuclear segmentation (*74*), and LaRoME plugin (<https://github.com/BIOP>) for labelling ROIs. To quantify the number of positive cells or the percentage of positive area, a threshold was set according to background signal. For the correlation analysis of the signal intensity of connexin 43 with respect to cTEVs positivity, and phospho-ERK1/2 with respect to vimentin positivity, an iterative study of the images was carried out consisting of: (i) establishing a grid of areas with a size of 5 or 50 µm^2^, for vimentin or connexin 43 studies, respectively; (2) measuring the average signal intensity of cTEVs in each area; (3) establishing cTEVs positive and negative areas considering background signal; (4) and, finally, measuring the signal intensities for connexin 43 or vimentin in the positive and negative areas. The quantification was performed in 3-4 randomly selected optical fields per section. At least two non-consecutive sections per specimen were used.

Cell apoptosis detection by TUNEL assay

Cell apoptosis was detected by means of the terminal deoxynucleotidyl transferase-mediated deoxyuridine triphosphate nick end labeling (TUNEL) assay. A commercial kit (Roche, Switzerland) was employed on myocardial sections following the manufacturer’s instructions. After the TUNEL assay, sections were stained with 0.5 μg/mL DAPI to verify the nuclear location of the TUNEL signals. Sections were observed with a Nikon A1R confocal microscope (Nikon, Japan).

Cell necrosis detection by IgG uptake assay

Cell necrosis was detected by means of IgG cardiomyocyte uptake assay (*75*). In brief, an anti-mouse IgG antibody developed in goat (Sigma-Aldrich, United States) was incubated overnight in histological sections overnight (1:500 dilution). The sections were also incubated in 1.0 mg/mL WGA labelled with Alexa Fluor (Thermo Fischer Scientific, United States) diluted 1:250 in PBS for labelling of cardiomyocyte cell membranes. Finally, sections were stained with DAPI. The sections were observed with a Nikon A1R confocal microscope (Nikon, Japan). The intracellular signal intensity was quantified as indicated for immunofluorescence techniques.

Interstitial collagen quantification and detection of organized collagen

For quantification of interstitial collagen, 8-µm histological sections were stained with picrosirius red and scanned with a Zeiss Axioscan 7 Scanner Microscope. Interstitial collagen organization was measured in the ventricular myocardium by second-harmonic generation (SHG) detection as described (*76*). In brief, organized collagen visualization was achieved using multiphoton SHG microscopy on a Zeiss LSM 780 Upright confocal and multiphoton microscope using a water-dipping Plan-Apochromat 40x/NA 1,0 objective (Zeiss, Germany). The tissue was excited at 860 nm and SHG signal was detected at 430 nm (10 nm window). Quantification was performed as described above in four randomly selected optical fields per section. At least two non-consecutive sections per specimen were used.

Neonatal cardiomyocyte immunofluorescence analysis

Cardiomyocytes were fixed with PFA4% at room temperature for 10 min, permeabilized with 0.2% Triton X-100 (Sigma-Aldrich) in 1% BSA (Sigma-Aldrich) for 5 min and blocked in SB for 1 hour at room temperature. Cells were incubated with primary antibody at 4°C overnight, and after washing, a secondary antibody was added for 1 hour at room temperature. All primary antibodies (Table S1) were diluted 1:100. Fluoromont G (BioNova, 0100-01) was used as mounting medium and allowed to cure overnight. Images were taken using a confocal microscope Zeiss LSM700 with Plan-Apochromat 40x/1.3 Oil DIC M27 objective. For image analysis, Fiji and a custom script written in ImageJ macro language were used. The outer membrane of cells was manually segmented and sarcomere markers were used to quantify number of cardiomyocytes. To average Troponin C intensity profiles, intensity values were first interpolated using cubic splines.

Neonatal cardiomyocyte dissociation assay

Cardiomyocytes were seeded in 8-well cell culture slides (MatTek, CCS-8). Cells were washed with HBSS and incubated with dissociation buffer (2.5 U/mL Dispase II in HBSS) (Sigma-Aldrich, D4693) at 37 °C till detachment of the cell monolayer from the bottom of the well. After detachment, monolayers were mechanically stressed by pipetting and the total number of resulting fragments was determined using a binocular stereo microscope (Leica, MZ FLIII). Images were acquired with a digital camera (Olympus, DP71). Fragments were counted with ImageJ-FIJI software.

Extraction of cell nuclei

To isolate cell nuclei, we followed the protocol described in (*77*). Briefly, tissues from three animals per group were resuspended in lysis buffer and homogenized using a Dounce homogenizer and a T10 UltraTurrax probe homogenizer. The homogenate was then filtered through a series of cell strainers (subsequent passages on 100, 70 and 40 μm strainers), and centrifuged at 1,000 g for 10 minutes to pellet the nuclei, which were washed twice with nuclei storage buffer (2.5M Sucrose, 1M Tris-HCl, pH = 7.2, 1M KCl, 1M MgCl_2_, 2M spermidine, protease inhibitor cocktail set III-EDTA-free, 40 units/mL RNase OUT) and resuspended in 2% BSA for subsequent purification by cell sorting. We enriched the sample in diploid and tetraploid nuclei based on their DAPI staining intensity in a FACSAria cell sorter (BD Biosciences). Purified nuclei were then processed for sequencing by CNIC’s Genomics Unit using 10X Genomics platform.

Single-nuclei RNA analysis

Cellranger (v6.1.1) pipeline from 10X Genomics was used to align sequencing data against refdata-gex-mm10-2020-A reference with two additional sequences, eGFP-uTEV3 and HaloTag-TEVs (*26*). Nuclei were filtered and clustered using Scater (v1.26.1) (*78*) and Seurat (v4.40) (*79*) R packages. Nuclei were filtered using a sequencing depth between 400 and 20,000, a minimum of 200 detected genes, a mitochondrial content below 5%, a cell fraction (proportion of unique molecular identifiers or UMIs the cell contributes to the median of UMIs per cell in the sample) above 0.2, a gene expression complexity (percentage of reads from the top 50 genes) below 60% and a hemoglobin gene set expression below 0.1%. eGFP-uTEV3, HaloTag-TEVs, mitochondrial and hemoglobin genes were excluded for downstream analysis. At the end of the filtering process, a total of 2,085 nuclei were retained. Doublets were identified using scDblFinder (v1.12.0) (*80*). Counts were normalized using default Seurat parameters, and the most variable 2,000 genes were identified using variance stabilizing transformation. Clustering at 0.75 resolution and dimensionality reduction of these nuclei were performed using 15 principal components. Cluster markers and differential expression analysis between conditions were performed using the MAST algorithm, testing only genes detected in more than 10% of nuclei. Clusters were manually annotated following the information in (*81*). Enrichment analysis was performed using PANTHER 19.0 (Gene Ontology). We used bar plots analysis considering DEG without fold-change thresholds to represent the top 10 enriched Cellular Component (CC), Biological Process (BP) and Molecular Function (MF) GO-Terms.

The analysis of expression dynamics in single-nuclei RNA sequencing data (i.e. RNA velocity) was performed using Velocyto (*33*), a package that allows estimating RNA velocities distinguishing between spliced and unspliced mRNAs in standard single-cell RNA sequencing protocols. Velocyto was then executed with default parameters and the refdata-gex-mm10-2020-A reference genome. After concatenation of the spliced and unspliced data from all experiments, results were merged with the outputs from single cell analyses performed with Seurat in R, and scVelo (*82*) was used for further processing. Preprocessing included gene selection by detection (the minimum number of both unspliced and spliced counts was set to 30), and by variability (keep 2,000 highly variable genes), normalization, and log1p transformation. First and second order moments were computed among the 30 nearest neighbors in the PCA space using 30 components. Cell-based RNA velocities were estimated by modeling the transcriptional dynamics of splicing kinetics using the standard model available in scVelo. Finally, these velocities were projected onto the previously computed UMAP and visualized as velocity vector fields.

Echocardiography

Animals were anesthetized using 1%-2% isoflurane in 100% oxygen, and non-invasive transthoracic echocardiography measurements were taken by personnel blinded to the study, using a high-resolution ultrasound imaging system with a 30-MHz transducer (Vevo 2100, MS400, VisualSonics, Canada). Standard parasternal long or short-axis views in two-dimensional (2D) mode and M-mode images of the left ventricle (LV) were acquired. M-mode images at the level of the papillary muscles were used to estimate LV end-diastolic and end-systolic volumes (LV Vol;d and LV Vol;s, respectively) and to calculate ejection fraction ((LV Vol;d – LV Vol;s) / LV Vol;d x 100). Diastolic function was evaluated by pulsed-wave Doppler using a 2D apical view to estimate mitral valve inflow.

Surface electrocardiographic recording

Animals were anesthetized using isoflurane inhalation (1.5%-2% volume in oxygen). Four-lead surface electrocardiograms were recorded for 1 minute using subcutaneous limb electrodes connected to an MP36R amplifier unit (BIOPAC Systems). Data acquisition and analysis were performed using AcqKnowledge (BIOPAC Systems). QTc intervals were estimated using the following expression (*83, 84*):

$QTc= \frac{QT}{\sqrt{RR/100}}$	(Equation 1)

Statistical analysis

In all figures, measurements are reported as mean ± standard error of the mean (SEM). The number of independent experiments is specified in the figure legends. Statistical significance was evaluated in GraphPad Prism 10. Unless stated otherwise, t-test was used to assess differences between groups. When experimental data did not follow a normal distribution, Mann-Whitney test was applied instead. Differences were considered statistically significant at *p < 0.05, **p < 0.01, ***p < 0.001, and ****p < 0.0001.

Supplementary Text S1. Production of a vector to drive GFP-uTEV3 expression under a cardiac troponin T promoter.

We used MluI and HindIII restriction enzymes to exchange promoters driving expression of GFP-uTEV3. With this aim, the following sequence for cardiac troponin T promoter including restriction sites was ordered as a G-block to IDT:

acgcgttgtacagcagtctgggctttcacaagacagcatttggggctgcggcagagggtcgggtccgaagcgctgccttatcagcgtccccagccctgggaggtgacaaaaggctggcttgtgtcagcccctcgggcactcacgtatctccatccgacgggtttaaaatagcaaaactctgaggccacacaatagcttgggcttatatgggctcctgtgggggaagggggagcacggagggggccggggccgctgctgccaaaatagcagctcacaagtgttgcattcctctctgggcgccgggcacattcctgctggctctgcccgccccggggtgggcgccggggggaccttaaagcctctgccccccaaggagcccttcccagacagccgccggcacccaccgctccgtgggacctctctggctaactagagaacccactgcttactggcttatcgaaattaatacgactcactatagggagacccaagctt

Figure S1. Supporting data on the expression of GFP-TEVp using AAVMYO infection.

(**A**) Detection of GFP-positive cells using immunofluorescence. (**B**) Evaluation of apoptosis in the myocardium of experimental groups using TUNEL (N = 5 animals per group). TUNEL-positive cells are indicated by red arrowheads. (**C**) IgG uptake evaluation to quantify necrosis (N = 5 animals per group). (**D**) Principal component analysis of bulk RNAseq data (N = 4 animals per group). (**E**) Quantification of the area occupied by collagen in the myocardium of wild-type (WT) animals injected with PBS or AAVMYO using Picrosirius red (N = 9-10 per group). All scale bars = 50 µm. Results are expressed as mean ± standard error of the mean (SEM). p-value <0.01(**), <0.001 (***), ). non-significant (ns). WGA: wheat germ agglutinin.

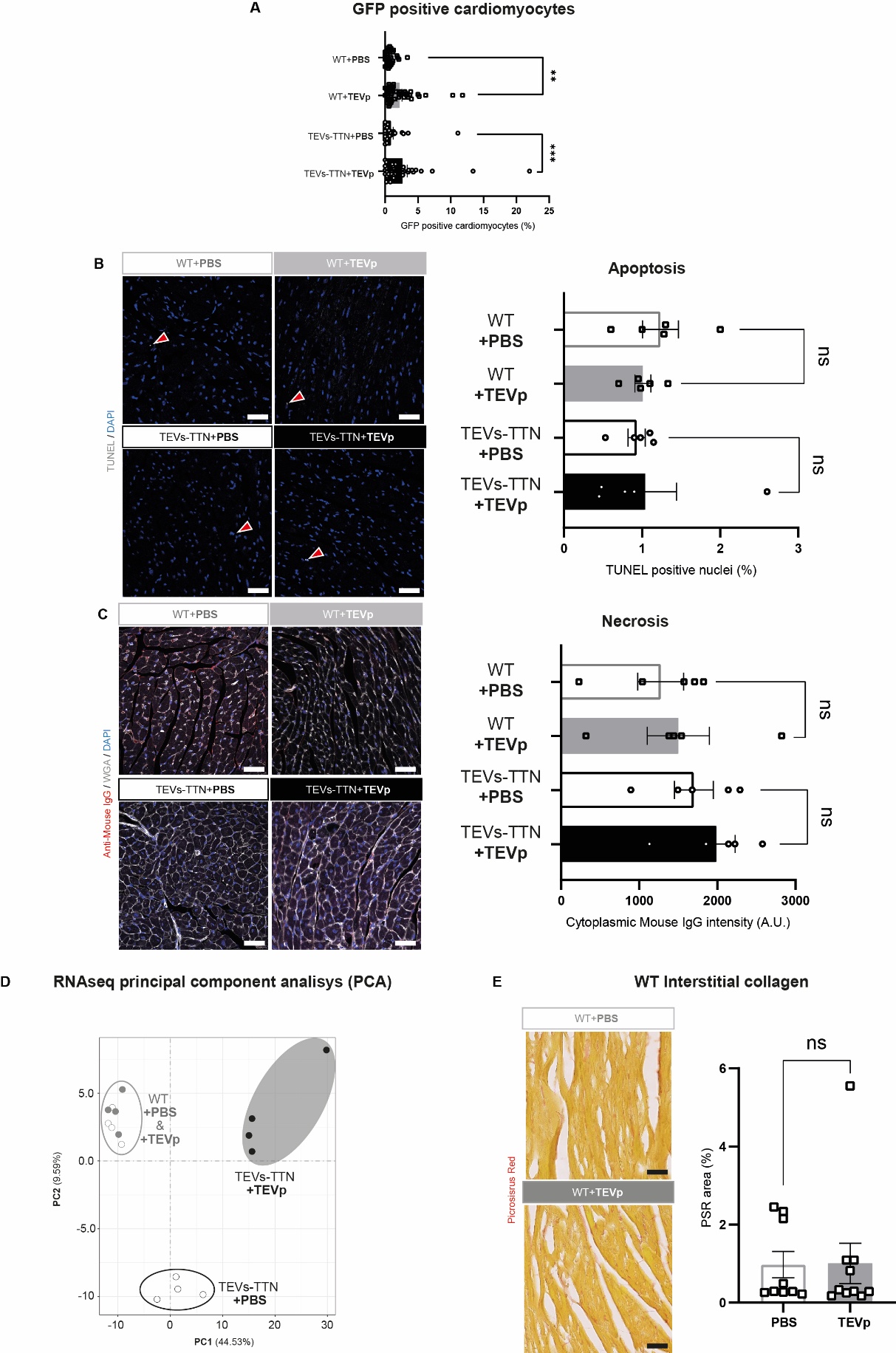

Figure S2. Experiments at higher doses of AAVMYO or using myocardium-specific cTnT promoter.

(**A**) Picrosirius red staining of collagen in the myocardium of mice injected with PBS or 3.82 x 10^13^ TEVp-expressing AAVMYO (N = 4 -9 animals per group) at 3 dpi. Control data already presented in **Figure 1**. (**B**) Scheme of the construct cTnT-GFP-TEVp used for specific expression in cardiomyocytes. (**C**) SDS-PAGE analysis of the fraction of full-length cardiac titin. (**D**) Assessment of the number of affected cardiomyocytes in ventricular myocardium of TEVs-TTN animals injected or not with AAVMYO expressing TEVp under the control of the cTnT promoter by their positivity to the antibody against cleaved TEV site (cTEVs, red) (N = 5 animals per group). (**E**) Picrosirius red staining of collagen in the myocardium of control TEVs-TTN mice (PBS) and those injected with AAVMYO expressing GFP-TEVp under the control of the cTnT promoter (N = 5 animals per group). All scale bars = 50 µm. Results are expressed as mean ± standard error of the mean (SEM). p-value < 0.05 (*), <0.01(**), <0.0001 (****). WGA: wheat germ agglutinin

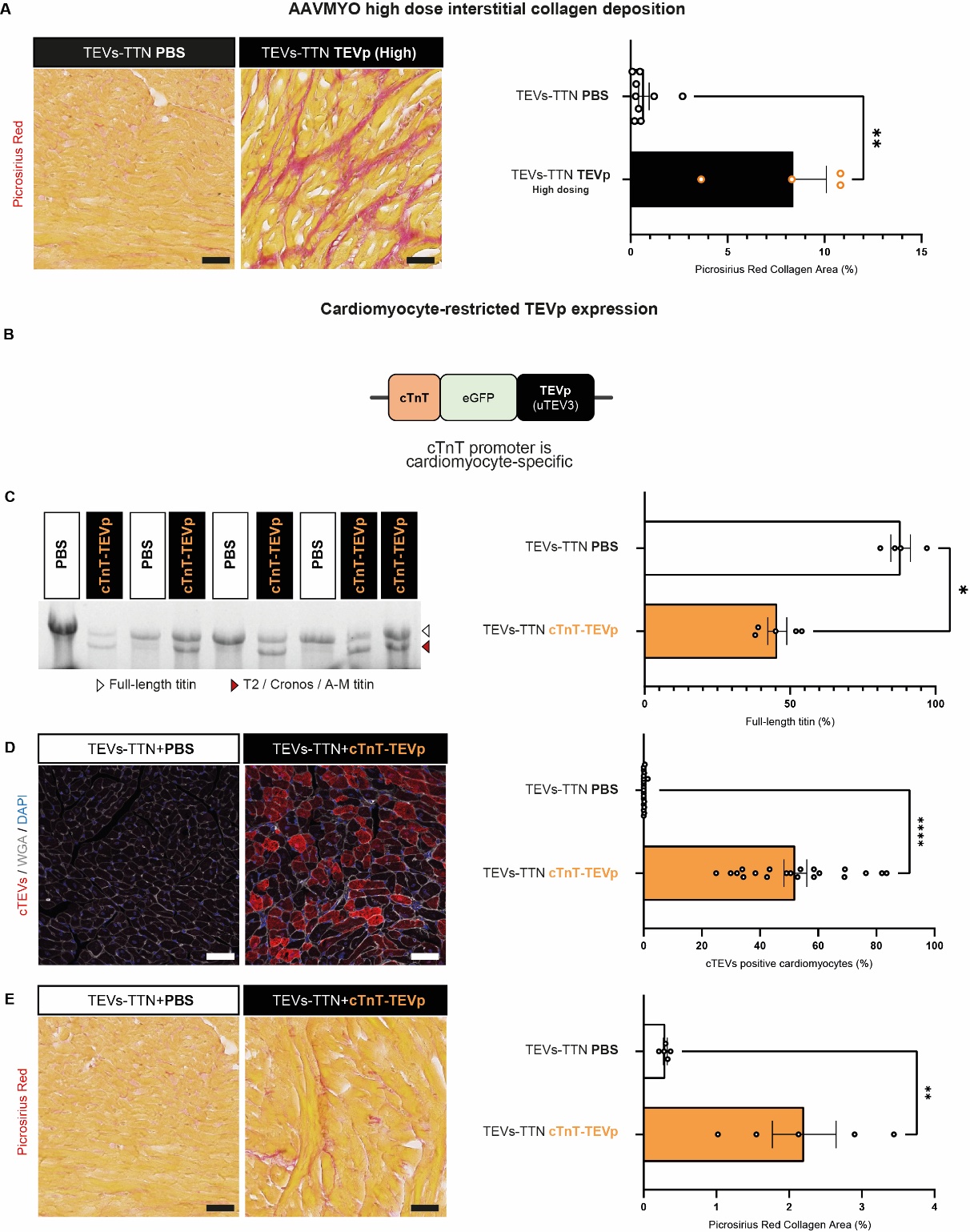

Figure S3. State of cardiac fibroblasts in wild-type animals upon expression of TEVp.

(**A-D**) Immunolocalization of vimentin (green)-positive fibroblasts expressing alpha-smooth muscle actin (red, αSMA) in WT mice injected or not with TEVp-expressing AAVMYO. (**E**) Quantification of results (N = 5 animals per group). (**F-I**) Immunolocalization and quantification of vimentin-positive fibroblasts expressing Ki67 (red) in WT mice injected or not with TEVp-expressing AAVMYO. (**J**) Quantification of results (N = 5 animals per group). All scale bars = 50 µm. Results are expressed as mean ± standard error of the mean (SEM). ). Non-significant p-value (ns). WGA: Wheat Germ Agglutinin. PC: pericardial cavity; LVL: left ventricle lumen.

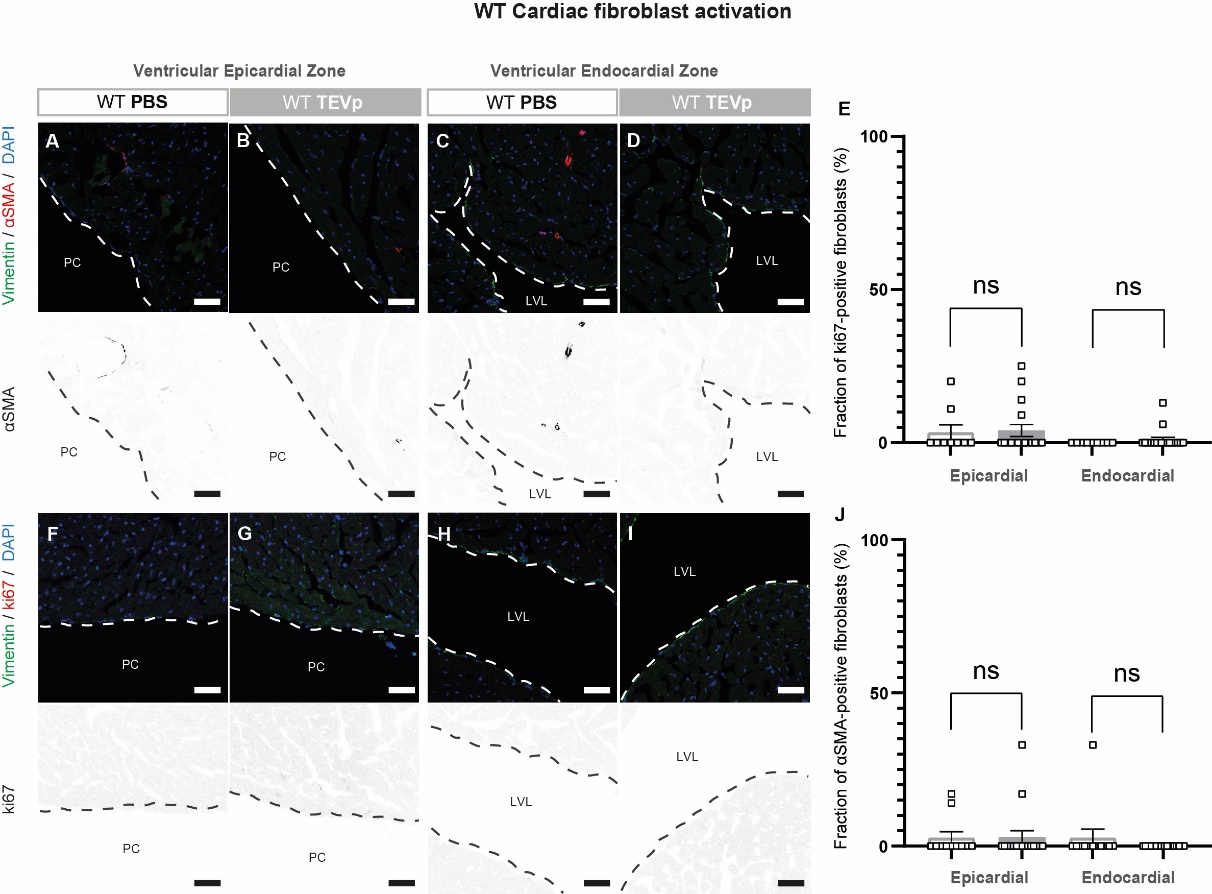

Figure S4. Evaluation of fibrotic pathways.

(**A**) Immunodetection of phospho-SMAD2/3 in histological sections from the myocardium of TEVs-TTN mice injected with PBS or TEVp-expressing AAVMYO (N = 7 animals per group). (**B**) Gene set enrichment analysis (GSEA) of the TGF-β signaling pathway using bulk RNAseq data from the myocardium of TEVs-TTN animals injected with PBS or TEVp-expressing AAVMYO (N = 4 animals per group; nominal p-value is indicated). (**C**) Immunodetection of phospho-p38 in the myocardium of TEVs-TTN animals injected with PBS or TEVp-expressing AAVMYO (N = 8 animals per group). (**D**) GSEA of the KRAS signaling pathway using bulk RNAseq data from the myocardium of TEVs-TTN animals injected with PBS or TEVp-expressing AAVMYO (N = 4 animals per group; nominal p-value is indicated). All scale bars = 50 µm. Results are expressed as mean ± standard error of the mean (SEM). ). Non-significant p-value (ns). WGA: wheat germ agglutinin.

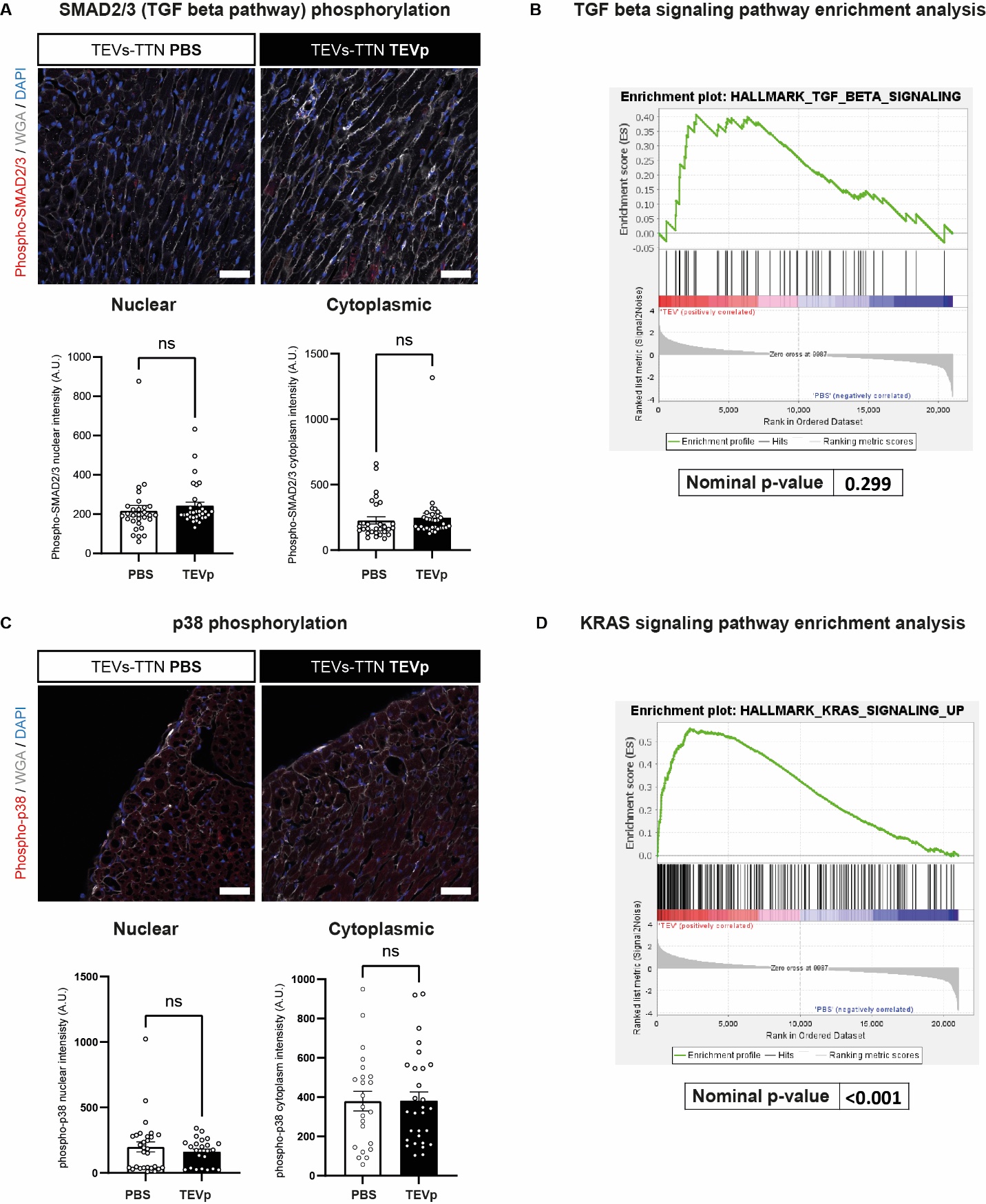

Figure S5. Identification of cell types in snRNAseq data.

(**A**) Uniform Manifold Approximation and Projection (UMAP) of the transcriptional state of nuclei from ventricular myocardial samples from the TEVs-TTN experimental groups indicating cell type annotation. (**B**) Cardiomyocyte identification by expression levels of TNNT2. (**C-F**) Top expressed gene in each annotated cell type. (**G**) Fraction of cell types represented in both experimental groups.

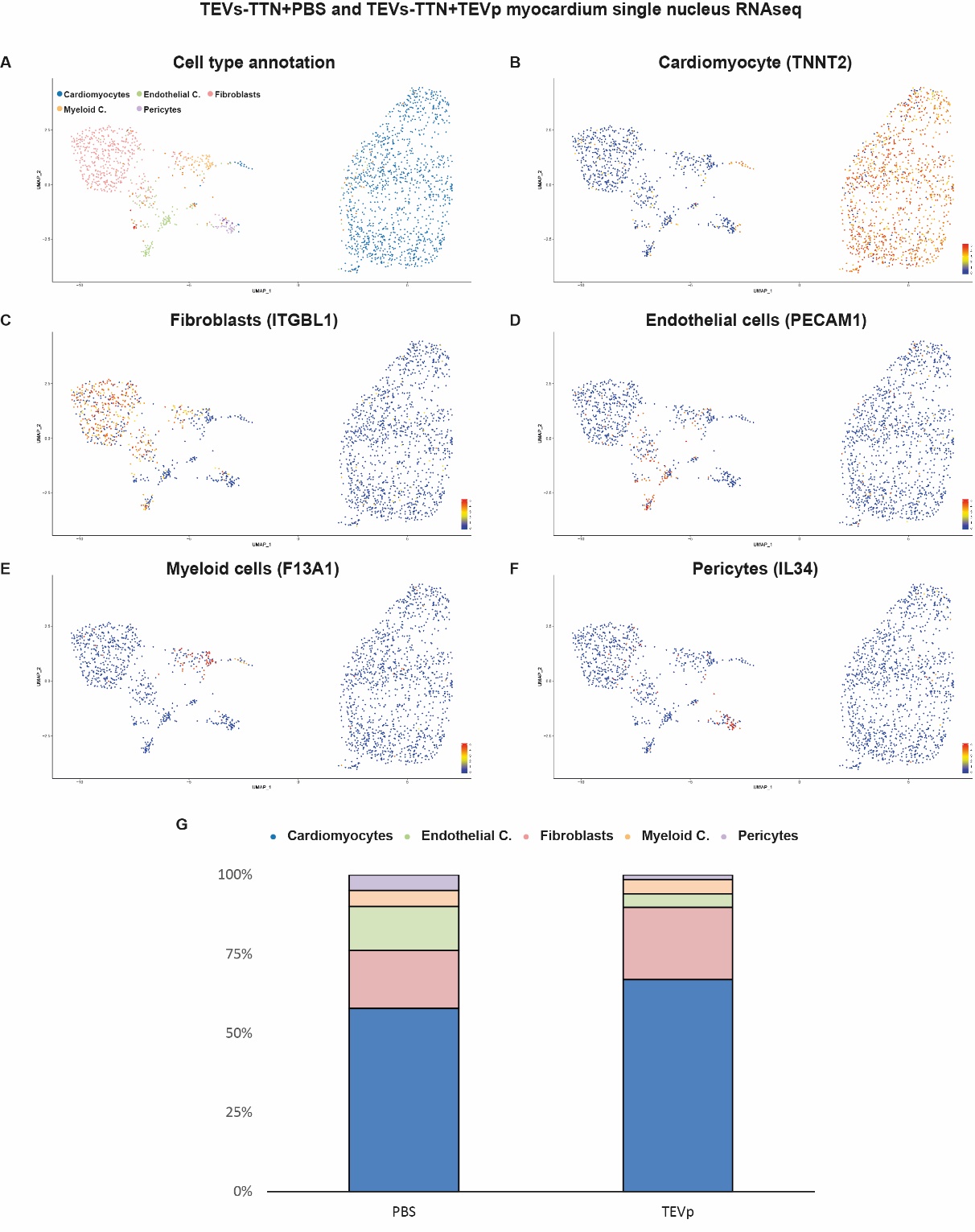

Figure S6. Early effects of titin cleavage three days after inoculation of a standard dose of TEVp-expressing AAVMYO.

(**A**) SDS-PAGE analysis of the fraction of full-length cardiac titin (**B**) Assessment of the number of affected cardiomyocytes in ventricular myocardium of TEVs-TTN animals injected or not with AAVMYO expressing TEVp by their positivity to the antibody against cTEVs (red). (**C**) Immunodetection of phospho-ERK1/2. (**D**) Immunolocalization and quantification of vimentin-positive fibroblasts expressing ki67 (red, arrowheads). (**E**) Picrosirius red staining of collagen in the myocardium of TEVs-TTN animals injected or not with TEVp-expressing AAVMYO. (**F**) Immunolocalization and quantification of vimentin (green)-positive fibroblasts expressing alpha-smooth muscle actin (red, αSMA). (**G**) Immunostaining of connexin 43. (**H**) Immunolocalization of integrin α5 – β1 complex. All scale bars = 50 µm. Results are expressed as mean ± standard error of the mean (SEM). N = 5 animals per group. p-value <0.001 (***), <0.0001 (****), non-significant (ns). WGA: wheat germ agglutinin. PC: pericardial cavity.

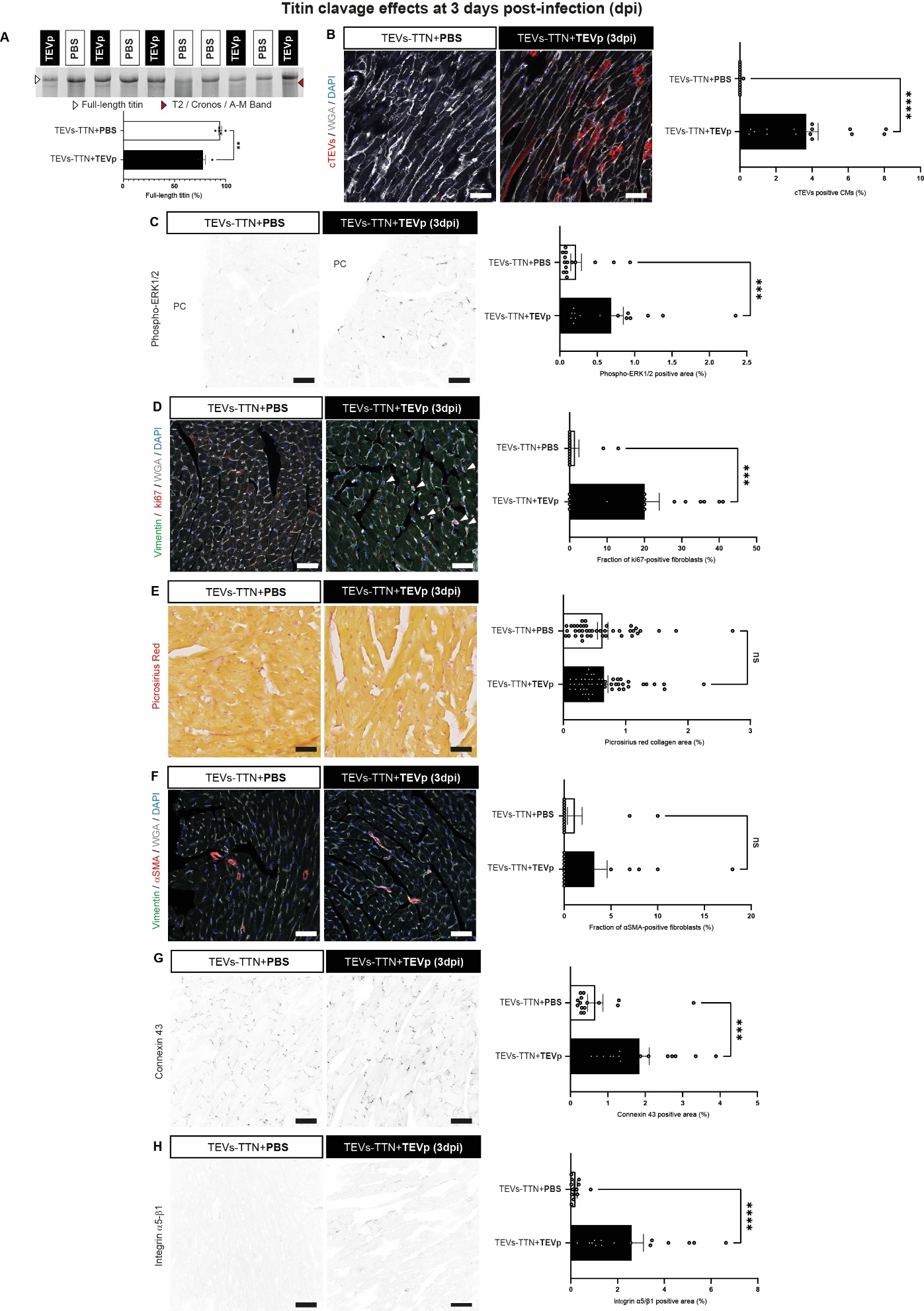

Figure S7. Electrocardiography study of mice subjected to titin cleavage at 5-6 dpi.

(A) Representative electrocardiograms indicating increased QRS and QT times in the TEVp-expressing TEVs-TTN mice with respect to PBS-injected controls. Bars indicate duration of QT interval from the second heartbeat in the recordings. (B-E) Quantification of electrophysiological parameters. N = 8 and 7 animals for PBS- and AAVMYO-injected groups, respectively. Results are expressed as mean ± standard error of the mean (SEM). p-value < 0.05 (*), <0.0001 (****).

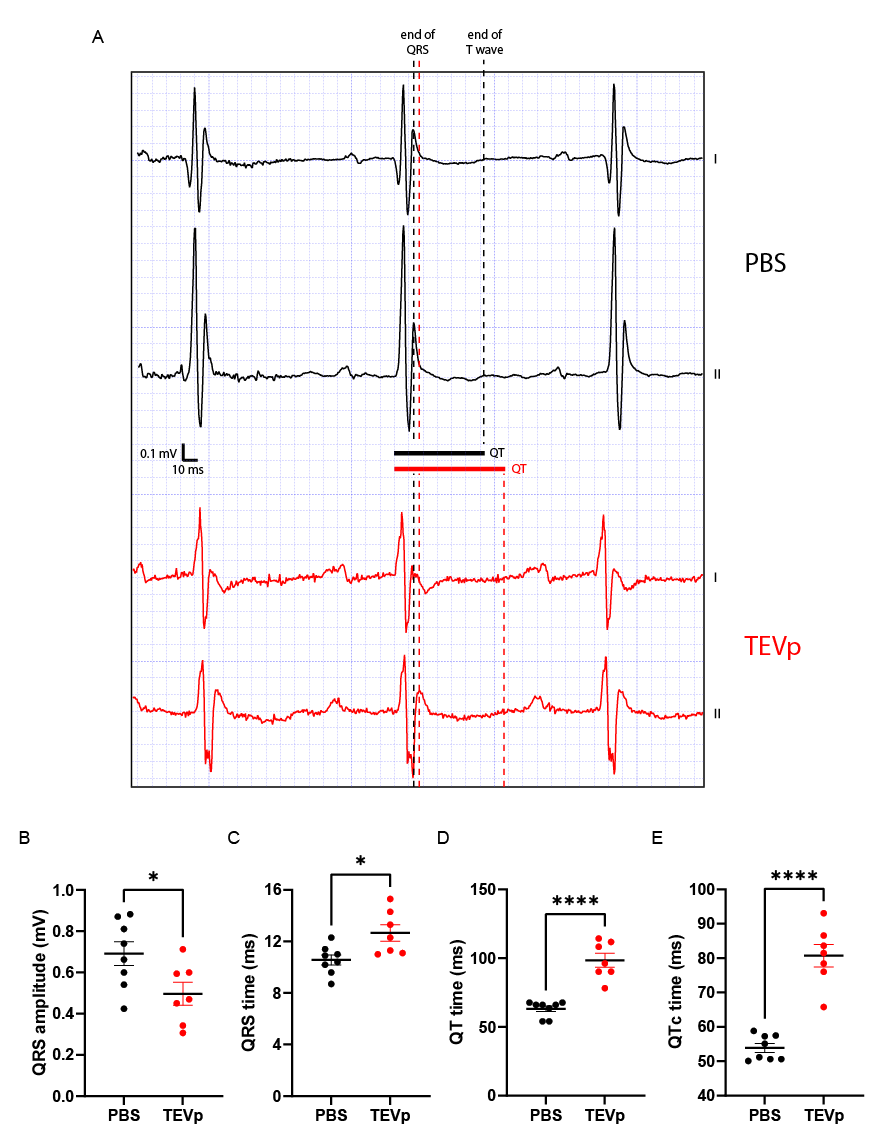

Figure S8. Echocardiographic study of cardiac function in mice subjected to titin cleavage at 5 dpi.

(**A-E**) Echocardiography functional parameters at 5 dpi of WT or TEVs-TTN mice, injected with TEVp-expressing AAVMYO or PBS (N = 5 animals per group). (**F**) Survival curve of a cohort of WT and TEVs-TTN mice injected with TEVp-expressing AAVMYO (N = 5 animals per group). QT time in the AAVMYO-injected TEVs-TTN mice is prolonged above reference values (84). Results are expressed as mean ± standard error of the mean (SEM). p–-value < 0.05 (*), non-significant (ns). Scheme adapted from ww.biorender.com.

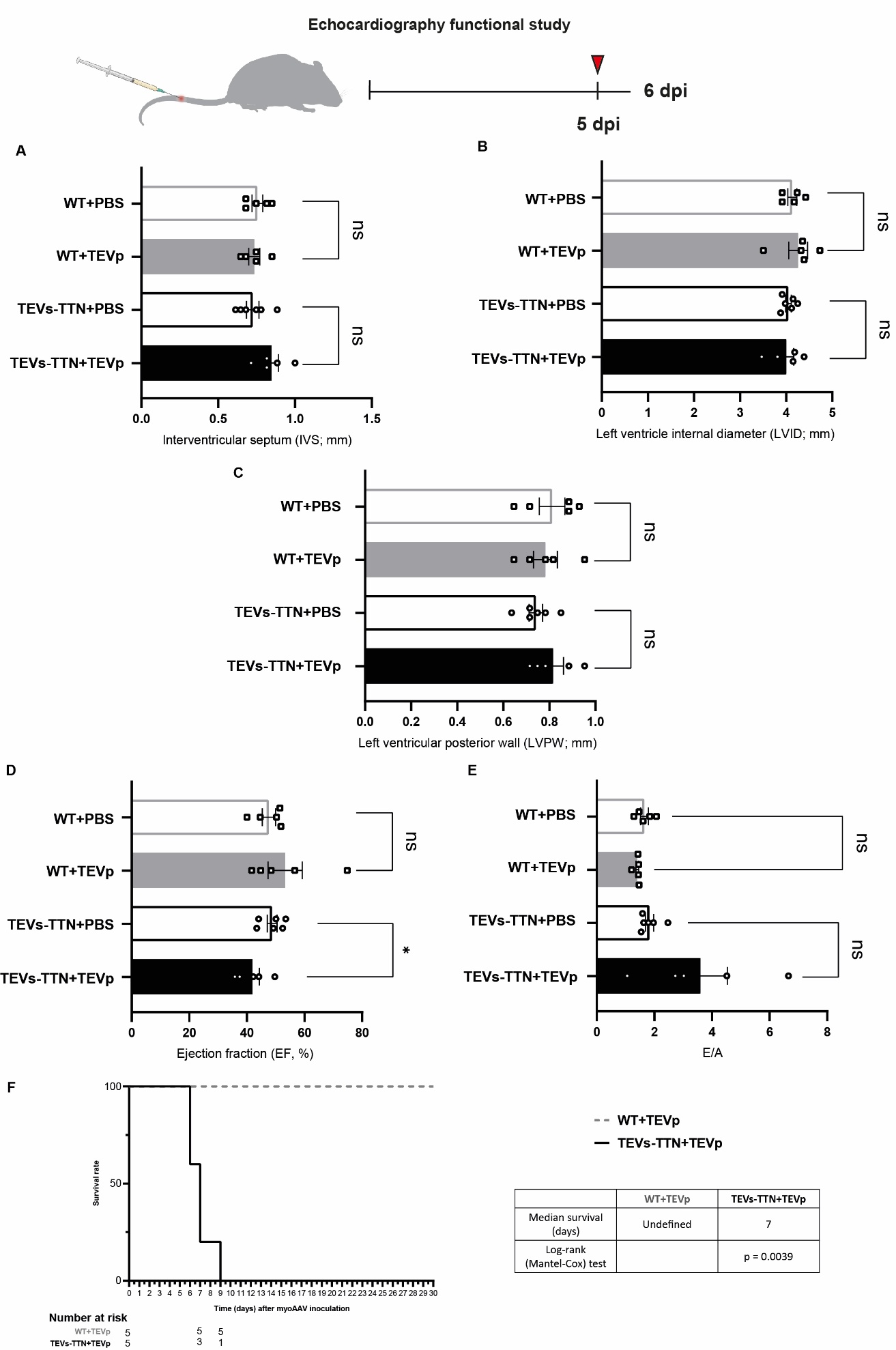

Figure S9. Quantification of full-length titin in human right ventricle from pulmonary hypertension (PAH) patients. Data from Figure 7B in S. Rain, M. L. Handoko, P. Trip, C. T.-J. Gan, N. Westerhof, G. J. Stienen, . . . F. S. de Man, Right Ventricular Diastolic Impairment in Patients With Pulmonary Arterial Hypertension. Circulation 128, 2016-2025 (2013). Results are expressed as mean ± standard error of the mean (SEM). p–-value < 0.05 (*).

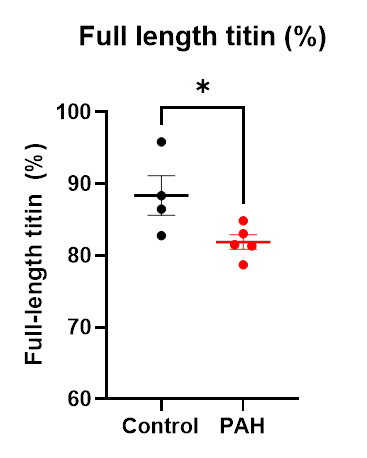

Table S1.

AAVMYO viral particles injected doses *in vivo*. See **Data S1** for individual values in relation to specific experimental cohorts.

| **Dose** | **Promoter** | **Mean injected**  **Viral Particles** |
| --- | --- | --- |
| *CMV-Standard* | CMV | 2.09 x 10^12^ |
| *CMV-High* | CMV | 3.02 x 10^12^ |
| *cTnT-Standard* | cTnT | 1.53 x 10^12^ |

Table S2.

Primary antibodies used for immunofluorescence.

| **Experiment** | **Target** | **Clone** | **Host** | **Reference** | **RRID** | **Dilution** |
| --- | --- | --- | --- | --- | --- | --- |
| *In vivo* | GFP | Polyclonal | Chicken | A10262, ThermoFisher Scientific | AB_2534023 | 1:100 |
|  | TEV cleavage site | Polyclonal | Rabbit | NBP2-37831, Novus Biologicals | AB_3297347 | 1:100 |
|  | Connexin 43 | Polyclonal | Goat | LS‑B9770, LSBio | N/A | 1:100 |
|  | Integrin α5-β1 | Monoclonal | Mouse | MAB2514, Sigma-Aldrich | AB_94626 | 1:100 |
|  | Vimentin | Polyclonal | Rabbit | ab45939, Abcam | AB_2257290 | 1:100 |
|  | Vimentin | Monoclonal | Mouse | V6630; Sigma-Aldrich | AB_477627 | 1:100 |
|  | α smooth muscle actin - Cy3 | Monoclonal | Mouse | C6198, Sigma-Aldrich | AB_476856 | 1:100 |
|  | ki67-PE | Monoclonal | Rat | 12-5698-82, ThermoFisher Scientific | AB_11150954 | 1:100 |
|  | Phospho-SMAD2 (Ser465/467)/SMAD3 (Ser423/425) | Monoclonal | Rabbit | 8828, Cell Signaling Technology | AB_2631089 | 1:100 |
|  | Phospho-p38 MAPK (Thr180/Tyr182) | Polyclonal | Rabbit | 9211, Cell Signaling Technology | AB_331641 | 1:100 |
|  | Phospho-p44/42 MAPK (Erk1/2) (Thr202/Tyr204) | Polyclonal | Rabbit | 9101, Cell Signaling Technology | AB_331646 | 1:100 |
| *In vitro* | Troponin T - Alexa Fluor 488 | Polyclonal | Rabbit | BS-10648R-A488, Bioss | N/A | 1:100 |
|  | Connexin 43 | Polyclonal | Rabbit | C6219, Merck | AB_476857 | 1:100 |

Data S1. (separate file)

AAV batch used per experiment and number of independent experiments replicated in laboratory.

Data S2. (separate file)

Newly generated custom computer code for image quantification.
