## Supplementary material for "Titin cleavage is a driver of cardiomyocyte disengagement and reactive myocardial fibrosis": DATA S2

**Image J (FIJI) Macro 1# Measure Area-Signal**

//CNIC Microscopy Unit 2023

//Detects Tissue area and measures Area, % of Area and Mean Intensity signal in Tissue, in up to 2 channels

macro "Analyze_Intensity_InTissue"{

//Getting Fiji and ROI Manager ready

run("Clear Results");

run("Close All");

run("Set Measurements...", "area display redirect=None decimal=3");

roiManager("Reset");

roiManager("Associate", "false");

roiManager("Centered", "false");

roiManager("UseNames", "true");

run("Line Width...", "line=1");

run("Colors...", "foreground=white background=black selection=yellow");

run("Options...", "iterations=1 count=1 black do=Nothing");

run("Bio-Formats Macro Extensions");

rowposition=0;

if(isOpen("Log")){

selectWindow("Log");

run("Close");

}

//Dialog Box

tissuethresholditems=newArray("Fixed Threshold Range", "AutoThreshold Method");

thresholdmethods=newArray("None","Default","Huang","Intermodes","IsoData","IJ_IsoData","Li","MaxEntropy","Mean","MinError","Minimum","Moments","Otsu","Percentile","RenyiEntropy","Shanbhag","Triangle","Yen");

IMGFormats=newArray(".lif", ".tif", ".TIFF", ".lsm", ".nd2", ".czi");

Dialog.create("Analysis Parameters");

Dialog.addChoice("Select image format", IMGFormats, ".nd2");

Dialog.addMessage("- Tissue ROI detection:", 14, "#1e00b9");

Dialog.addRadioButtonGroup("", tissuethresholditems, 1, 2, "Fixed Threshold Range");

Dialog.addMessage("Based on your previous selection, set parameters:", 12, "#0036ff");

Dialog.addNumber("Set min Threshold value:", 35);

Dialog.addChoice("Select Autothreshold Method:", thresholdmethods, "Huang");

Dialog.addCheckbox("Threshold visual inspection?", false);

Dialog.addMessage("	");

Dialog.addMessage("- Channel Signal detection:", 14, "#1e00b9");

Dialog.addNumber("Channel with signal to measure", 2);

Dialog.addToSameRow();

Dialog.addString("Name of signal to measure", "COL1");

Dialog.addRadioButtonGroup("", tissuethresholditems, 1, 2, "AutoThreshold Method");

Dialog.addMessage("Based on your previous selection, set parameters:", 12, "#0036ff");

Dialog.addNumber("Set min Threshold value:", 40);

Dialog.addChoice("Select Autothreshold Method:", thresholdmethods, "Triangle");

Dialog.addCheckbox("Threshold visual inspection?", false);

Dialog.addMessage("	");

Dialog.addMessage("- Extra Channel Signal detection:", 14, "#1e00b9");

Dialog.addCheckbox("Do you want to analyze another channel?", true);

Dialog.addMessage("Please specify:", 12, "#0036ff");

Dialog.addNumber("2nd Channel with signal to measure", 4);

Dialog.addToSameRow();

Dialog.addString("Name of 2nd signal to measure", "SHG");

Dialog.addRadioButtonGroup("", tissuethresholditems, 1, 2, "AutoThreshold Method");

Dialog.addMessage("Based on your previous selection, set parameters:", 12, "#0036ff");

Dialog.addNumber("Set min Threshold value:", 40);

Dialog.addChoice("Autothreshold method to detect 2nd signal", thresholdmethods, "Triangle");

Dialog.addCheckbox("Threshold visual inspection?", false);

Dialog.show();

//Get dialog box info

ImageFormat=Dialog.getChoice();

TissueThresholdMethod=Dialog.getRadioButton();

TissueFixedThresholdMethod=Dialog.getNumber();

TissueAutoThresholdMethod=Dialog.getChoice();

TissueVisualThreshold=Dialog.getCheckbox();

SignalChannel=Dialog.getNumber();

SignalLabel=Dialog.getString();

SignalThresholdMethod=Dialog.getRadioButton();

SignalFixedThresholdMethod=Dialog.getNumber();

SignalAutoThresholdMethod=Dialog.getChoice();

SignalVisualThreshold=Dialog.getCheckbox();

Extrachannel=Dialog.getCheckbox();

ExtraSignalChannel=Dialog.getNumber();

ExtraSignalLabel=Dialog.getString();

ExtraSignalThresholdMethod=Dialog.getRadioButton();

ExtraSignalFixedThresholdMethod=Dialog.getNumber();

ExtraSignalAutoThresholdMethod=Dialog.getChoice();

ExtraSignalVisualThreshold=Dialog.getCheckbox();

//Print Dialog Box options

print("=====================================================================================");

print("Analysis Parameters:");

print("--------------------");

print("Method to detect Tissue is:"+TissueThresholdMethod);

if(TissueThresholdMethod=="Fixed Threshold Range"){print("Tissue min threshold is: "+TissueFixedThresholdMethod);

}else{print("Tissue Auto threshold method is: "+TissueAutoThresholdMethod);}

print("Signal to measure is "+SignalLabel+" in channel "+SignalChannel);

print("Method to detect Signal is:"+SignalThresholdMethod);

if(SignalThresholdMethod=="Fixed Threshold Range"){print("Signal min threshold is: "+SignalFixedThresholdMethod);

}else{print("Signal Auto threshold method is: "+SignalAutoThresholdMethod);}

if(Extrachannel==1) {

print("Extra Signal to measure is "+ExtraSignalLabel+" in channel "+ExtraSignalChannel);

if(ExtraSignalThresholdMethod=="Fixed Threshold Range"){print("Extra Signal min threshold is: "+ExtraSignalFixedThresholdMethod);

}else{print("Extra Signal Auto threshold method is: "+ExtraSignalAutoThresholdMethod);}

}

print("=====================================================================================");

//----

//Loop to process all files in a folder.

dir = getDirectory("Choose folder with files to process...");

getDateAndTime(year, month, dayOfWeek, dayOfMonth, hour, minute, second, msec);

output = dir+"Results_Macro_Run#_"+year+"_"+month+"_"+dayOfMonth+"_"+hour+"_"+minute+"_"+second+File.separator;

lista = getFileList(dir);

File.makeDirectory(output);

print("=======================");

print("Macro Analysis has started around "+hour+":"+minute+":"+second);

for (i=0; i<lista.length; i++) {

if (endsWith(lista[i], ""+ImageFormat+"")==1) {

Ext.setId(dir+lista[i]);

Ext.getCurrentFile(archivo);

Ext.getSeriesCount(series);

for (s=0; s<series; s++) {

Ext.setSeries(s);

Ext.getSizeC(sizeC);

Ext.getSeriesName(imagen_actual);

if(sizeC>1){

run("Bio-Formats Importer", "open=["+archivo+"] autoscale color_mode=Composite view=Hyperstack stack_order=XYCZT series_"+s+1);

setBatchMode(false);

flag=0;

myimagename = getTitle();

myimagename=replace(myimagename, "/", "_");

print("=======================");

print("Beginning analysis of image "+myimagename);

FLUOImg=getImageID();

rename("ORI");

Stack.getDimensions(widthFLUOImg, heightFLUOImg, channelsFLUOImg, slicesFLUOImg, framesFLUOImg);

getVoxelSize(widthORI, heightORI, depthORI, unitORI);

if(bitDepth==8){upper=255;}else if(bitDepth==16){upper=65535;}

setBatchMode(true);

//Detect Tissue

for(ch=1;ch<=channelsFLUOImg;ch++){

Stack.setChannel(ch);

run("Enhance Contrast", "saturated=0.35");

}

run("RGB Color");

RGB=getImageID();

run("8-bit");

//run("Gaussian Blur...", "sigma="+heightORI+" scaled");

run("Median...", "radius=1");

if(TissueThresholdMethod=="Fixed Threshold Range"){

setThreshold(TissueFixedThresholdMethod, upper);

getThreshold(myfinaltissuethresholdMIN, myfinaltissuethresholdMAX);

}else{

setAutoThreshold(TissueAutoThresholdMethod+" dark");

getThreshold(myfinaltissuethresholdMIN, myfinaltissuethresholdMAX);

}

//run("Analyze Particles...", "size=100-Infinity show=Masks");

//setThreshold(1, 255, "raw");

if(TissueVisualThreshold==1){

print("Tissue visual threshold adjustment selected");

selectImage(RGB);

rename("TISSUE");

setBatchMode("show");

run("Threshold...");

waitForUser("Tissue Visual Threshold Adjustment", "Adjust Tissue threshold\n \nPress OK when done to continue...");

getThreshold(myfinaltissuethresholdMIN, myfinaltissuethresholdMAX);

selectWindow("TISSUE");

setBatchMode("hide");

}

selectImage(RGB);

rename("TISSUE");

List.setMeasurements;

myAreafractionTissue=List.getValue("%Area");

if(myAreafractionTissue>0){

run("Create Selection");

roiManager("add");

tissueroiindex=roiManager("count")-1;

roiManager("select",tissueroiindex);

run("Enlarge...", "enlarge=-4 pixel");

run("Enlarge...", "enlarge=4 pixel");

roiManager("update");

roiManager("rename", "Tissue");

List.setMeasurements;

AreaTissue=List.getValue("Area");

}else{

selectImage(FLUOImg);

setBatchMode("show");

//Count ROIs

AllROIs=roiManager("count");

do {

waitForUser("Tissue Area Not Detected--Starting Manual Selection", "Draw tissue area and use *t* key to add it to ROI Manager\n \n When done, press OK to continue with analysis");

AllROIs=roiManager("count");

}while (AllROIs==0);

tissueroiindex=roiManager("count")-1;

roiManager("select",tissueroiindex);

roiManager("rename", "Tissue");

List.setMeasurements;

AreaTissue=List.getValue("Area");

selectImage(FLUOImg);

setBatchMode("hide");

}

//close("Mask of ORI (RGB)");

close("TISSUE");

for(ch=1;ch<=channelsFLUOImg;ch++){

Stack.setChannel(ch);

resetMinAndMax();

}

//Detect and measure signal using macro function MeasureMyChannel

if(SignalChannel<=channelsFLUOImg){

myDataforChannel=MeasureMyChannel (FLUOImg, SignalChannel, SignalLabel);

ChannelArea=myDataforChannel[0];

ChannelAreaPercentage=myDataforChannel[1];

ChannelMean=myDataforChannel[2];

minThresholdSignal=myDataforChannel[3];

maxThresholdSignal=myDataforChannel[4];

}else{

Dialog.create("Image does not have enough channels...");

Dialog.addNumber("Please set again Channel with signal to measure", 2);

Dialog.show();

SignalChannel=Dialog.getNumber();

print("Redefined Signal to measure is "+SignalLabel+" in channel "+SignalChannel);

myDataforChannel=MeasureMyChannel (FLUOImg, SignalChannel, SignalLabel);

ChannelArea=myDataforChannel[0];

ChannelAreaPercentage=myDataforChannel[1];

ChannelMean=myDataforChannel[2];

minThresholdSignal=myDataforChannel[3];

maxThresholdSignal=myDataforChannel[4];

}

if(Extrachannel==1) {

if(ExtraSignalChannel<=channelsFLUOImg){

myDataforExtraChannel=MeasureMyChannel(FLUOImg, ExtraSignalChannel, ExtraSignalLabel);

ExtraChannelArea=myDataforExtraChannel[0];

ExtraChannelAreaPercentage=myDataforExtraChannel[1];

ExtraChannelMean=myDataforExtraChannel[2];

ExtraminThresholdSignal=myDataforExtraChannel[3];

ExtramaxThresholdSignal=myDataforExtraChannel[4];

}else{

Dialog.create("Image does not have enough channels...");

Dialog.addNumber("Please set again Extra Channel with signal to measure", 2);

Dialog.show();

ExtraSignalChannel=Dialog.getNumber();

print("Redefined Extra Signal to measure is "+ExtraSignalLabel+" in channel "+ExtraSignalChannel);

myDataforExtraChannel=MeasureMyChannel(FLUOImg, ExtraSignalChannel, ExtraSignalLabel);

ExtraChannelArea=myDataforExtraChannel[0];

ExtraChannelAreaPercentage=myDataforExtraChannel[1];

ExtraChannelMean=myDataforExtraChannel[2];

ExtraminThresholdSignal=myDataforExtraChannel[3];

ExtramaxThresholdSignal=myDataforExtraChannel[4];

}

}

//Results table

run("Clear Results");

if (isOpen("TempResults")){

Table.rename("TempResults", "Results");

}

setResult("Image", rowposition,myimagename);

setResult("Area Units (^2)", rowposition, unitORI);

setResult("Tissue Area", rowposition, AreaTissue);

setResult("Ch "+SignalChannel+" " +SignalLabel+" Area", rowposition, ChannelArea);

setResult("% "+SignalLabel+" in Tissue Area", rowposition, ChannelAreaPercentage);

setResult(SignalLabel+" Mean intensity", rowposition, ChannelMean);

if(Extrachannel==1) {

setResult("Ch "+ExtraSignalChannel+" " +ExtraSignalLabel+" Area", rowposition, ExtraChannelArea);

setResult("% "+ExtraSignalLabel+" in Tissue Area", rowposition, ExtraChannelAreaPercentage);

setResult(ExtraSignalLabel+" Mean intensity", rowposition, ExtraChannelMean);

setResult(ExtraSignalLabel+"Threshold Range", rowposition, "["+ExtraminThresholdSignal+" ,"+ ExtramaxThresholdSignal+"]");

}

setResult("Tissue Threshold Range", rowposition, "["+myfinaltissuethresholdMIN+" ,"+ myfinaltissuethresholdMAX+"]");

setResult(SignalLabel+"Threshold Range", rowposition, "["+minThresholdSignal+" ,"+ maxThresholdSignal+"]");

updateResults();

rowposition++;

saveAs("Results", output+"Results.xls");

saveAs("Text", output+"Results.txt");

if (isOpen("Results")){

Table.rename("Results", "TempResults");

}

run("Close All");

//Save ROI manager items

roiManager("deselect");

allmyroielements=roiManager("count");

if(allmyroielements!=0){

roiManager("save", output+myimagename+"-ROIs.zip");

roiManager("reset");

}else{

print("No ROI elements detected for image " + myimagename);

roiManager("reset");

}

if(isOpen("Log")){

selectWindow("Log");

saveAs("Text", output+"Log");

}

}

}

}

}

getDateAndTime(yearEND, monthEND, dayOfWeekEND, dayOfMonthEND, hourEND, minuteEND, secondEND, msecEND);

print("Macro Analysis done at "+hourEND+":"+minuteEND+":"+secondEND);

print("=================THE=END================");

if(isOpen("Log")){

selectWindow("Log");

saveAs("Text", output+"Log");

}

if(isOpen("Log")){

selectWindow("Log");

run("Close");

}

waitForUser("Analysis done, please take a look to your results:\n \n " + output);

}

//------------------------------------------------------------------

//Macro function:

function MeasureMyChannel (inputID, channelinput, ROIlabel){

selectImage(inputID);

run("Select None");

selectImage(inputID);

run("Duplicate...", "title="+ROIlabel+" duplicate channels="+channelinput);

selectWindow(ROIlabel);

roiManager("select",tissueroiindex);

setBackgroundColor(0, 0, 0);

run("Clear Outside");

if(SignalChannel==channelinput){

if(SignalThresholdMethod=="Fixed Threshold Range"){

setThreshold(SignalFixedThresholdMethod, upper);

getThreshold(myfinalthresholdMIN, myfinalthresholdMAX);

}else{

setAutoThreshold(SignalAutoThresholdMethod+" dark");

getThreshold(myfinalthresholdMIN, myfinalthresholdMAX);

}

if(SignalVisualThreshold==1){

print("Visual threshold adjustment selected for channel "+channelinput);

selectWindow(ROIlabel);

setBatchMode("show");

run("Threshold...");

waitForUser("Visual Threshold Adjustment for channel "+channelinput+" ("+ROIlabel+")", "Adjust threshold. Press OK when done to continue...");

getThreshold(myfinalthresholdMIN, myfinalthresholdMAX);

selectWindow(ROIlabel);

setBatchMode("hide");

}

}else if (ExtraSignalChannel==channelinput){

if(ExtraSignalThresholdMethod=="Fixed Threshold Range"){

setThreshold(ExtraSignalFixedThresholdMethod, upper);

getThreshold(ExtramyfinalthresholdMIN, ExtramyfinalthresholdMAX);

}else{

setAutoThreshold(ExtraSignalAutoThresholdMethod+" dark");

getThreshold(ExtramyfinalthresholdMIN, ExtramyfinalthresholdMAX);

}

if(ExtraSignalVisualThreshold==1){

print("Visual threshold adjustment selected for channel "+channelinput);

selectWindow(ROIlabel);

setBatchMode("show");

run("Threshold...");

waitForUser("Visual Threshold Adjustment for channel "+channelinput+" ("+ROIlabel+")", "Adjust threshold. Press OK when done to continue...");

getThreshold(ExtramyfinalthresholdMIN, ExtramyfinalthresholdMAX);

selectWindow(ROIlabel);

setBatchMode("hide");

}

}

mydatainarray=newArray(5);

selectWindow(ROIlabel);

List.setMeasurements("limit");

mydatainarray[0]=List.getValue("Area");

mydatainarray[1]=List.getValue("%Area");

mydatainarray[2]=List.getValue("Mean");

if((Extrachannel==1) && (ExtraSignalChannel==channelinput)){

mydatainarray[3]=ExtramyfinalthresholdMIN;

mydatainarray[4]=ExtramyfinalthresholdMAX;

}else if((Extrachannel==1) && (ExtraSignalChannel!=channelinput)){

mydatainarray[3]=myfinalthresholdMIN;

mydatainarray[4]=myfinalthresholdMAX;

}else if(SignalChannel==channelinput){

mydatainarray[3]=myfinalthresholdMIN;

mydatainarray[4]=myfinalthresholdMAX;

}

if(mydatainarray[0]>0){

run("Create Selection");

if(selectionType() !=0){

roiManager("add");

signalroiindex=roiManager("count")-1;

roiManager("select",signalroiindex);

roiManager("rename", ROIlabel);

}

}else {print("No "+ROIlabel+ " detected for image channel "+ channelinput);}

return mydatainarray;

close(ROIlabel);

}

**Image J (FIJI) Macro21# Heatmaps – Colocalization**

//CNIC Microscopy Unit 2024

macro "The_HeatMaper"{

/*

//HeatMap Legend

HM1=ChA

HM2=ChB

HM3=HM1*HM2

HM4=HM2-HM3

HM5=HM1-HM3

*/

//1. Create Dialog Box

FijiLUTs=newArray("Red Hot", "Cyan Hot", "Yellow Hot", "Magenta Hot", "Green", "Red", "Blue", "Magenta", "Cyan", "Yellow", "Grays");

IMGFormats=newArray(".lif", ".tif", ".TIFF", ".lsm", ".nd2", ".czi");

Dialog.create("Analysis Parameters");

Dialog.addChoice("Select image format", IMGFormats, ".nd2");

Dialog.addNumber("Specify ROIs size in microns", 10);

Dialog.addMessage("HeatMaps Channel Info:", 12, "#0036ff");

Dialog.addMessage("	");

Dialog.addNumber("Channel A number", 2);

Dialog.addString("Channel A Label", "Titina");

Dialog.addNumber("Channel A min Threshold", 650);

Dialog.addChoice("HM1 Channel A LUT", FijiLUTs, "Red Hot");

Dialog.addMessage("	");

Dialog.addNumber("Channel B number", 3);

Dialog.addString("Channel B Label", "Vinculina");

Dialog.addNumber("Channel B min Threshold", 402);

Dialog.addChoice("HM2 Channel B LUT", FijiLUTs, "Cyan Hot");

Dialog.addMessage("Choose HeatMap Operations LUTs:", 12, "#0036ff");

Dialog.addMessage("	");

Dialog.addChoice("HM3 (HM1*HM2) LUT", FijiLUTs, "Yellow Hot");

Dialog.addChoice("HM4 (HM2-HM3) LUT", FijiLUTs, "Magenta Hot");

Dialog.addChoice("HM5 (HM1-HM3) LUT", FijiLUTs, "Green");

Dialog.show();

//2. Read Dialog Box Info

ImageFormat=Dialog.getChoice();

boxSide=Dialog.getNumber();

ChANumber=Dialog.getNumber();

ChALabel=Dialog.getString();

ChAMINThreshold=Dialog.getNumber();

ChALUT=Dialog.getChoice();

ChBNumber=Dialog.getNumber();

ChBLabel=Dialog.getString();

ChBMINThreshold=Dialog.getNumber();

ChBLUT=Dialog.getChoice();

ProjectionLUT=Dialog.getChoice();

HM4LUT=Dialog.getChoice();

HM5LUT=Dialog.getChoice();

//3. Getting Fiji and ROI Manager ready

setBatchMode(true);

rowposition=0;

run("Clear Results");

run("Close All");

roiManager("show all with labels");

run("Set Measurements...", " redirect=None decimal=6");

roiManager("Reset");

roiManager("Associate", "false");

roiManager("Centered", "false");

roiManager("UseNames", "true");

run("Line Width...", "line=0");

run("Colors...", "foreground=white background=black selection=yellow");

if(isOpen("Log")){

selectWindow("Log");

run("Close");

}

//4. Loop to process all files in a folder.

input=getDirectory("Choose folder with images to process...");

lista = getFileList(input);

getDateAndTime(year, month, dayOfWeek, dayOfMonth, hour, minute, second, msec);

output = input+"Results_Macro_Run#_"+year+"_"+month+"_"+dayOfMonth+"_"+hour+"_"+minute+"_"+second+File.separator;

File.makeDirectory(output);

run("Bio-Formats Macro Extensions");

for (ii=0; ii<lista.length; ii++) {

if (endsWith(lista[ii],ImageFormat)==1) {

Ext.setId(input+lista[ii]);

Ext.getCurrentFile(archivo);

Ext.getSeriesCount(series);

for (s=0; s<series; s++) {

Ext.setSeries(s);

Ext.getSeriesName(imagen_actual);

Ext.getSizeX(sizeX);

Ext.getSizeY(sizeY);

run("Bio-Formats Importer", "open=["+archivo+"] autoscale color_mode=Composite view=Hyperstack stack_order=XYCZT series_"+s+1);

myimagename=getTitle();

rename("ORI");

myORIimage=getImageID();

if(bitDepth()=="8"){

maxValue=255;

}else if((bitDepth()=="16")&&(ImageFormat==".nd2")){

maxValue=4095;

}

getDimensions(widthORI, heightORI, channelsORI, slicesORI, framesORI);

getVoxelSize(widthPX, heightPX, depthPX, unitPX);

boxSidepixels=boxSide/widthPX;

//5. Select Image area

selectImage(myORIimage);

run("Select All");

roiManager("add");

lastItem=roiManager("count")-1;

roiManager("select", lastItem);

roiManager("rename", "AnalizedArea");

//6. Split Image Area into defined size boxes

roiManager("select", 0);

getSelectionBounds(xROI, yROI, widthROI, heightROI);

getDimensions(widthImage, heightImage, channelsImage, slicesImage, framesImage);

areaROI = widthROI*heightROI;

getSelectionCoordinates(x, y);

boxArea= pow(boxSidepixels, 2);

numBoxY = floor(heightROI/boxSidepixels); // number of boxes that will fit along the height

numBoxX = floor(widthROI/boxSidepixels); // "" width

remainX = (widthROI - (numBoxX*boxSidepixels))/2; // remainder distance left when all boxes fit

remainY = (heightROI - (numBoxY*boxSidepixels))/2;

miniROIsX=newArray(numBoxX);

miniROIsY=newArray(numBoxY);

for (i=0; i<numBoxX; i++) {

miniROIsX[i]=(xROI+(boxSidepixels*i+1));

}

for (i=0; i<numBoxY; i++) {

miniROIsY[i]=(yROI+(boxSidepixels*i+1));

}

newImage("BIN", "8-bit black", widthImage, heightImage, 1);

roiManager("select", 0);

run("Colors...", "foreground=white background=black selection=yellow");

run("Fill", "slice");

for (i=0; i<miniROIsX.length; i++) {

for (j=0; j<miniROIsY.length; j++) {

selectWindow("BIN");

makeRectangle((remainX/numBoxX)+miniROIsX[i], (remainY/numBoxY)+miniROIsY[j], boxSidepixels, boxSidepixels);

List.setMeasurements;

BinArea=List.getValue("%Area");

if(BinArea>95){

roiManager("add");

}

}

}

//7. Create empty heatmap images

newImage(ChALabel, "32-bit black", widthImage, heightImage, 1);

setVoxelSize(widthPX, heightPX, depthPX, unitPX);

newImage(ChBLabel, "32-bit black", widthImage, heightImage, 1);

setVoxelSize(widthPX, heightPX, depthPX, unitPX);

newImage("HM3", "32-bit black", widthImage, heightImage, 1);

setVoxelSize(widthPX, heightPX, depthPX, unitPX);

newImage("HM4", "32-bit black", widthImage, heightImage, 1);

setVoxelSize(widthPX, heightPX, depthPX, unitPX);

newImage("HM5", "32-bit black", widthImage, heightImage, 1);

setVoxelSize(widthPX, heightPX, depthPX, unitPX);

//8. Remove background intensities by image thresholding

selectWindow("ORI");

run("32-bit");

Stack.setChannel(ChANumber);

setThreshold(ChAMINThreshold, maxValue);

run("Create Selection");

if(selectionType()!=-1){

setBackgroundColor(0, 0, 0);

run("Clear Outside", "slice");

run("Select None");

}else{

run("Select All");

setBackgroundColor(0, 0, 0);

run("Clear", "slice");

run("Select None");

print("No signal above threshold found for ch:"+ChANumber+" in image "+myimagename);

}

Stack.setChannel(ChBNumber);

setThreshold(ChBMINThreshold, maxValue);

run("Create Selection");

if(selectionType()!=-1){

setBackgroundColor(0, 0, 0);

run("Clear Outside", "slice");

run("Select None");

}else{

run("Select All");

setBackgroundColor(0, 0, 0);

run("Clear", "slice");

run("Select None");

print("No signal above threshold found for ch:"+ChBNumber+" in image "+myimagename);

}

//9. Measure Mean Intensity in both channels

Stack.setDisplayMode("color");

allmyrois=roiManager("count");

myRoi=newArray(allmyrois-1);

myRedMeanIntensity=newArray(allmyrois-1);

myGreenMeanIntensity=newArray(allmyrois-1);

for(j=1;j<allmyrois;j++){

selectWindow("ORI");

roiManager("select", j);

roiManager("rename", "ROI-"+j);

myroiname=Roi.getName;

myRoi[j-1]=myroiname;

roiManager("select", j);

Stack.setChannel(ChANumber);

List.setMeasurements;

myRedMeanIntensity[j-1]=List.getValue("Mean");

Stack.setChannel(ChBNumber);

List.setMeasurements;

myGreenMeanIntensity[j-1]=List.getValue("Mean");

}

close("BIN");

//10. Normalize HeatMaps Intensity (max value will be set as 1; if normalization gives NaN, then equal value to 0)

//Array.show("Raw Data", myGreenMeanIntensity, myRedMeanIntensity);

Array.getStatistics(myGreenMeanIntensity, minGreen, maxGreen, meanGreen, stdDevGreen);

Array.getStatistics(myRedMeanIntensity, minRed, maxRed, meanRed, stdDevRed);

myRedMeanIntensityNormalized=newArray(allmyrois-1);

myGreenMeanIntensityNormalized=newArray(allmyrois-1);

for(j=0;j<myGreenMeanIntensity.length;j++){

myRedMeanIntensityNormalized[j]=myRedMeanIntensity[j]/maxRed;

if(isNaN(myRedMeanIntensityNormalized[j])){myRedMeanIntensityNormalized[j]=0;}

myGreenMeanIntensityNormalized[j]=myGreenMeanIntensity[j]/maxGreen;

if(isNaN(myGreenMeanIntensityNormalized[j])){myGreenMeanIntensityNormalized[j]=0;}

}

//Array.show("Normalized Data", myGreenMeanIntensityNormalized, myRedMeanIntensityNormalized);

//11. Assign normalized values to Heatmap images

for(j=1;j<allmyrois;j++){

selectWindow(ChALabel);

roiManager("select", j);

setForegroundColor(myRedMeanIntensityNormalized[j-1], myRedMeanIntensityNormalized[j-1], myRedMeanIntensityNormalized[j-1]);

run("Fill", "slice");

valuetochangeR=parseFloat(myRedMeanIntensityNormalized[j-1]);

changeValues(0, 255, valuetochangeR);

selectWindow(ChBLabel);

roiManager("select", j);

setForegroundColor(myGreenMeanIntensityNormalized[j-1], myGreenMeanIntensityNormalized[j-1], myGreenMeanIntensityNormalized[j-1]);

run("Fill", "slice");

valuetochangeG=parseFloat(myGreenMeanIntensityNormalized[j-1]);

changeValues(0, 255, valuetochangeG);

}

//12. Assign LUTs to Heatmaps

selectWindow(ChALabel);

run("Select None");

resetMinAndMax();

run(ChALUT);

saveAs("tif", output+myimagename+"_HM1_"+ChALabel);

rename("HM1_"+ChALabel);

selectWindow(ChBLabel);

run("Select None");

resetMinAndMax();

run(ChBLUT);

saveAs("tif", output+myimagename+"_HM2_"+ChBLabel);

rename("HM2_"+ChBLabel);

//13. Project one heatmap over the other (multiplication)

ProjectionImageNormalized=newArray(allmyrois-1);

for(j=0;j<myGreenMeanIntensity.length;j++){

ProjectionImageNormalized[j]=myRedMeanIntensityNormalized[j]*myGreenMeanIntensityNormalized[j];

}

//Asign values to Projection heatmap

for(j=1;j<allmyrois;j++){

selectWindow("HM3");

roiManager("select", j);

setForegroundColor(ProjectionImageNormalized[j-1], ProjectionImageNormalized[j-1], ProjectionImageNormalized[j-1]);

run("Fill", "slice");

valuetochange=parseFloat(ProjectionImageNormalized[j-1]);

changeValues(0, 255, valuetochange);

}

run(ProjectionLUT);

saveAs("tif", output+myimagename+"_HM3_Projection_HM1_"+ChALabel+"_x_HM2_"+ChBLabel);

rename("HM3_Projection_HM1_"+ChALabel+"_x_HM2_"+ChBLabel);

//14. Subtract projection to Channel B

SubtrationBNormalized=newArray(allmyrois-1);

for(j=0;j<myGreenMeanIntensity.length;j++){

SubtrationBNormalized[j]=myGreenMeanIntensityNormalized[j]-ProjectionImageNormalized[j];

}

//Asign values to SubtractB heatmap

for(j=1;j<allmyrois;j++){

selectWindow("HM4");

roiManager("select", j);

setForegroundColor(SubtrationBNormalized[j-1], SubtrationBNormalized[j-1], SubtrationBNormalized[j-1]);

run("Fill", "slice");

valuetochange=parseFloat(SubtrationBNormalized[j-1]);

changeValues(0, 255, valuetochange);

}

run(HM4LUT);

saveAs("tif", output+myimagename+"_HM4_Subtraction_HM2_"+ChBLabel+"_&_HM3_Projection");

rename("HM4_Subtraction_HM2_"+ChBLabel+"_&_HM3_Projection");

//15. Subtract projection to Channel A

SubtrationANormalized=newArray(allmyrois-1);

for(j=0;j<myGreenMeanIntensity.length;j++){

SubtrationANormalized[j]=myRedMeanIntensityNormalized[j]-ProjectionImageNormalized[j];

}

//Asign values to SubtractA heatmap

for(j=1;j<allmyrois;j++){

selectWindow("HM5");

roiManager("select", j);

setForegroundColor(SubtrationANormalized[j-1], SubtrationANormalized[j-1], SubtrationANormalized[j-1]);

run("Fill", "slice");

valuetochange=parseFloat(SubtrationANormalized[j-1]);

changeValues(0, 255, valuetochange);

}

run(HM5LUT);

saveAs("tif", output+myimagename+"_HM5_Subtraction_HM1_"+ChALabel+"_&_HM3_Projection");

rename("HM5_Subtraction_HM1_"+ChALabel+"_&_HM3_Projection");

//Array.show("DATA", myRoi, myRedMeanIntensity, myRedMeanIntensityNormalized, myGreenMeanIntensity, myGreenMeanIntensityNormalized, ProjectionImageNormalized,SubtrationANormalized,SubtrationBNormalized);

//16. Create Result table

run("Clear Results");

if (isOpen("TempResults")){

Table.rename("TempResults", "Results");

}

for(data=0;data<allmyrois-1;data++){

setResult("Image", rowposition, myimagename);

setResult("ROI #", rowposition, myRoi[data]);

setResult(ChALabel+" Raw Mean Intensity", rowposition, myRedMeanIntensity[data]);

setResult(ChBLabel+" Raw Mean Intensity", rowposition, myGreenMeanIntensity[data]);

setResult("HM1:"+ChALabel+" Normalized Mean Intensity", rowposition, myRedMeanIntensityNormalized[data]);

setResult("HM2:"+ChBLabel+" Normalized Mean Intensity", rowposition, myGreenMeanIntensityNormalized[data]);

setResult("HM3: Projection (HM1*HM2) Image Normalized Mean Intensity", rowposition, ProjectionImageNormalized[data]);

setResult("HM4: Subtration (HM2-HM3) Image Normalized Mean Intensity", rowposition, SubtrationBNormalized[data]);

setResult("HM5: Subtration (HM1-HM3) Image Normalized Mean Intensity", rowposition, SubtrationANormalized[data]);

updateResults();

rowposition++;

}

//17. Create HeatMaps montage:

selectWindow("ORI");

run("Make Composite");

if(channelsORI==4){

Stack.setChannel(1);

run("Blue");

Stack.setChannel(2);

run("Grays");

Stack.setChannel(3);

run("Red");

Stack.setChannel(4);

run("Green");

}else if (channelsORI==3){

Stack.setChannel(1);

run("Blue");

Stack.setChannel(2);

run("Green");

Stack.setChannel(3);

run("Red");

}else if (channelsORI==2){

Stack.setChannel(1);

run("Green");

Stack.setChannel(2);

run("Red");

}

run("RGB Color");

run("Scale Bar...", "width=20 height=20 horizontal bold");

rename("RGB-"+myimagename);

close("ORI");

//First, change all heatmaps to RGB

imagesopen=nImages;

for(img=1;img<=imagesopen;img++){

selectImage(img);

run("RGB Color");

}

//Then, create Montage RGB image

run("Images to Stack", "use keep");

run("Make Montage...", "columns=3 rows=2 scale=1 label");

saveAs("tif", output+myimagename+"_HeatMap-Montage");

//18. Save ROIs

roiManager("deselect");

roiManager("save", output+myimagename+"_ROIs.zip");

roiManager("reset");

//19. Add empty row on result table

setResult("Image", rowposition, "	");

setResult("ROI #", rowposition, "	");

setResult(ChALabel+" Raw Mean Intensity", rowposition, "	");

setResult(ChBLabel+" Raw Mean Intensity", rowposition, "	");

setResult("HM1:"+ChALabel+" Normalized Mean Intensity", rowposition, "	");

setResult("HM2:"+ChBLabel+" Normalized Mean Intensity", rowposition, "	");

setResult("HM3: Projection (HM1*HM2) Image Normalized Mean Intensity", rowposition, "	");

setResult("HM4: Subtration (HM2-HM3) Image Normalized Mean Intensity", rowposition, "	");

setResult("HM5: Subtration (HM1-HM3) Image Normalized Mean Intensity", rowposition, "	");

updateResults();

rowposition++;

//20. Save Results and prepare table for next round

selectWindow("Results");

saveAs("Results", output+"HeatMap-Results.xls");

selectWindow("Results");

saveAs("Text", output+"HeatMap-Results.txt");

if (isOpen("Results")){

Table.rename("Results", "TempResults");

}

//21. Close All

run("Close All");

}

}

}

waitForUser("Heatmaping is done, you can find your macro results here: \n "+ output);

}

**Image J (FIJI) Macro 3# Cells and nuclei intensity analysis**

//CNIC Microscopy Unit 2022

//This macro requires Cellpose installed in the computer: https://github.com/MouseLand/cellpose

//This macro requires Cellpose wrapper plugin:https://github.com/BIOP/ijl-utilities-wrappers

//This macro requires LaRoMe plugin:https://github.com/BIOP/ijp-LaRoMe

//This macro requires StarDist plugin:https://imagej.net/plugins/stardist

macro "Cells+Nuclei_Analysis"{

//Getting Fiji and ROI Manager ready

run("Clear Results");

roiManager("Reset");

roiManager("Associate", "false");

roiManager("Centered", "false");

roiManager("UseNames", "true");

run("Colors...", "foreground=white background=black selection=yellow");

dir = getDirectory("Select folder with files to analyze...");

getDateAndTime(year, month, dayOfWeek, dayOfMonth, hour, minute, second, msec);

results = dir+"Results"+"_"+hour+"h-"+minute+"min-"+second+"sec-"+msec+"msec"+File.separator;

File.makeDirectory(results);

lista = getFileList(dir);

//setBatchMode(true);

//Create Dialog box for macro parameters input

items=newArray("yeah", "No need");

Dialog.createNonBlocking("Please complete parameter-boxes before analysis");

Dialog.addString("Image Format	", ".nd2");

Dialog.addMessage("Image Channels info: \n ", 16, "#0066ff");

//--

Dialog.addNumber("Cell Detection Channel #	", 3);

Dialog.addToSameRow();

Dialog.addString("Set Channel Label	", "WGA");

Dialog.addToSameRow();

Dialog.addChoice("Do you want to perform Cell analysis?", items, "yeah");

Dialog.addToSameRow();

Dialog.addNumber("% of Max Signal for + Cells detection	", 45);

Dialog.addToSameRow();

Dialog.addNumber("Threshold Factor for + Cells detection (factor*Mean of Intensity means)	", 1.3);

//--

Dialog.addMessage("--------", 16, "#0066ff");

Dialog.addNumber("Nuclei Detection Channel #	", 1);

Dialog.addToSameRow();

Dialog.addString("Set Channel Label	", "DAPI");

Dialog.addToSameRow();

Dialog.addChoice("Do you want to perform Nuclei analysis?", items, "yeah");

Dialog.addToSameRow();

Dialog.addNumber("% of Max Signal for + Nuclei detection	", 60);

Dialog.addToSameRow();

Dialog.addNumber("Threshold Factor for + Nuclei detection (factor*Mean of Intensity means)	", 1.5);

//--

Dialog.addMessage("--------", 16, "#0066ff");

Dialog.addNumber("Signal to analize, Channel #	", 4);

Dialog.addToSameRow();

Dialog.addString("Set Channel Label	", "MyStuff");

Dialog.show();

//Get info from dialog

myimageformat=Dialog.getString();

Cellch=Dialog.getNumber();

CellChannelLabel=Dialog.getString();

CellAnalysis=Dialog.getChoice();

PercentageMaxSignalAnalysisCh=Dialog.getNumber();

PercentageDifferenceSignalCells=Dialog.getNumber();

Nucch=Dialog.getNumber();

NucChannelLabel=Dialog.getString();

NucleiAnalysis=Dialog.getChoice();

PercentageMaxSignalNucCh=Dialog.getNumber();

PercentageDifferenceSignalNuclei=Dialog.getNumber();

Analysisch=Dialog.getNumber();

AnalysisChannelLabel=Dialog.getString();

//Print dialog box options in Log window

print("------------------------------------------------");

print("------------------------------------------------");

print("Analysis run # "+hour+"h-"+minute+"min-"+second+"sec-"+msec+"msec");

print("------------------------------------------------");

print("Dialog Box options:");

print("------------------------------------------------");

print("Image info: Format="+myimageformat);

print("Channel for analysis # "+Analysisch+"; Channel label="+AnalysisChannelLabel);

if (CellAnalysis=="yeah"){

print("Cell analysis selected");

print("Cell Channel # "+Cellch+"; Channel label="+CellChannelLabel);

print("% of Max Signal for + Cells detection: "+PercentageMaxSignalAnalysisCh+" %");

print("Threshold Factor for + Cells detection: "+PercentageDifferenceSignalCells);

}

if (NucleiAnalysis=="yeah"){

print("Nuclei analysis selected");

print("Nuclei Channel # "+Nucch+"; Channel label="+NucChannelLabel);

print("% of Max Signal for + Nuclei detection: "+PercentageMaxSignalNucCh+" %");

print("Threshold Factor for + Nuclei detection: "+PercentageDifferenceSignalNuclei);

}

print("------------------------------------------------------------------------------------------------------------------------------------");

run("Bio-Formats Macro Extensions");

myposition=0;

for (i=0; i<lista.length; i++) {

if (endsWith(lista[i],myimageformat)==1) {

Ext.setId(dir+lista[i]);

Ext.getCurrentFile(archivo);

Ext.getSeriesCount(series);

for (s=0; s<series; s++) {

Ext.setSeries(s);

run("Bio-Formats Importer", "open=["+archivo+"] autoscale color_mode=Composite view=Hyperstack stack_order=XYCZT series_"+s+1);

myimagename=getTitle();

ORI=getImageID();

Stack.getDimensions(widthORI, heightORI, channelsORI, slicesORI, framesORI);

getVoxelSize(widthvoxel, heightvoxel, depthvoxel, unitvoxel);

print("------------------------------------------------");

print("Analyzing image "+	myimagename+ ". Area results will be expressed in ^2 "+ unitvoxel);

selectImage(ORI);

run("Duplicate...", "title="+AnalysisChannelLabel+" duplicate channels="+Analysisch);

ANALYSIS=getImageID();

if (CellAnalysis=="yeah"){

//****************Begining of CELLs analysis*********************************************************

//Cell channel image pre-processing

selectImage(ORI);

run("Duplicate...", "title="+CellChannelLabel+" duplicate channels="+Cellch);

CELLS=getImageID();

selectImage(CELLS);

run("Enhance Local Contrast (CLAHE)", "blocksize=127 histogram=256 maximum=3 mask=*None*");

run("Subtract Background...", "rolling=50");

run("Gaussian Blur...", "sigma=0.6");

//Cell detection using Cellpose

run("Cellpose Advanced", "diameter=60 cellproba_threshold=0.6 flow_threshold=10 anisotropy=1.0 diam_threshold=40.0 model=cyto2 nuclei_channel=0 cyto_channel=1 dimensionmode=2D stitch_threshold=-1.0 omni=false cluster=false additional_flags=0");

CellposeImg=getImageID();

rename("CELLP");

//Save cellpose label image

//selectImage(CellposeImg);

//saveAs(".tif", results+myimagename+"_CellposeLabelImg");

//Extract ROIs from Cellpose label image

run("Label image to ROIs", "");

//Rename final ROIs	using macro function

var allmyCELLrois=roiManager("count");

myCellsROINames=RENAMEmyROIs(0, allmyCELLrois, "C", "white");

roiManager("deselect");

//Measure Cell ROI parameters on channel of interest

run("Set Measurements...", "area mean display redirect=None decimal=3");

selectImage(ANALYSIS);

myCellsROIArea=MEASUREmyROIs(0, allmyCELLrois, "Area", 0);

myCellsROIMean=MEASUREmyROIs(0, allmyCELLrois, "Mean", 0);

//Create Cells total data Results table

Array.show("Cell_Data_Results", myCellsROINames, myCellsROIArea,myCellsROIMean);

selectWindow("Cell_Data_Results");

saveAs("Results", results+myimagename+"_Ch_"+AnalysisChannelLabel+"_Cell_Data_Results.xls");

selectWindow(myimagename+"_Ch_"+AnalysisChannelLabel+"_Cell_Data_Results.xls");

run("Close");

//Obtain threshold for positive signal CELL classification (% of Max Signal found will be used)

////Get array statistics to find max value

Array.getStatistics(myCellsROIMean, minMeanCellsValue, maxMeanCellsValue, meanMeanCellsValue);

print("Max mean value of all cells detected "+ maxMeanCellsValue);

print("Mean mean value of all cells detected "+ meanMeanCellsValue);

print("Min mean value of all cells detected "+ minMeanCellsValue);

ThresholdValueCells=(maxMeanCellsValue*(PercentageMaxSignalAnalysisCh/100));

DifferenceOfSignalsCells=meanMeanCellsValue*PercentageDifferenceSignalCells;

print("Threshold value used for positive cell classification on channel "+ AnalysisChannelLabel+" will be >= to "+ ThresholdValueCells);

print("Second Threshold value used for positive cell classification on channel "+ AnalysisChannelLabel+" will be > to "+ DifferenceOfSignalsCells);

//CELLs classification

////New array for positive/negative labeling of cells. Set counters

CELLPositiveORNegativeSignal=newArray(allmyCELLrois);

PosCell=0;

NegCell=0;

for(data=0;data<myCellsROIMean.length;data++){

if(myCellsROIMean[data]>=ThresholdValueCells && myCellsROIMean[data]>DifferenceOfSignalsCells){

CELLPositiveORNegativeSignal[data]="Positive";

PosCell++;

roiManager("select", data);

roiManager("rename", "+Cell_"+data+1);

roiManager("Set Color", "Green");

roiManager("Set Line Width", 1);

}else{

CELLPositiveORNegativeSignal[data]="Negative";

NegCell++;

roiManager("select", data);

roiManager("rename", "-Cell_"+data+1);

roiManager("Set Color", "Blue");

roiManager("Set Line Width", 1);

}

}

if(PosCell!=0){

//Create array with only data from positive cells detected

CELLPositiveMeanData=newArray(PosCell);

CELLPositiveAreaData=newArray(PosCell);

CELLPositiveROIName=newArray(PosCell);

positionarray=0;

for(data=0;data<myCellsROIMean.length;data++){

if(CELLPositiveORNegativeSignal[data]=="Positive"){

CELLPositiveMeanData[positionarray]=myCellsROIMean[data];

CELLPositiveAreaData[positionarray]=myCellsROIArea[data];

CELLPositiveROIName[positionarray]=myCellsROINames[data];

positionarray++;

}

}

//Create only positive cells data Results table

Array.show("Positive_Cells_Data_Results", CELLPositiveROIName, CELLPositiveAreaData,CELLPositiveMeanData);

selectWindow("Positive_Cells_Data_Results");

saveAs("Results", results+myimagename+"_Ch_"+AnalysisChannelLabel+"_Positive_Cells_Data_Results.xls");

selectWindow(myimagename+"_Ch_"+AnalysisChannelLabel+"_Positive_Cells_Data_Results.xls");

run("Close");

//Get statistics of only positive cells array

Array.getStatistics(CELLPositiveMeanData, dataIdonotneed, dataIdonotneed, meanMEANPositiveCells, stdDevMEANPositiveCells);

Array.getStatistics(CELLPositiveAreaData, dataIdonotneed, dataIdonotneed, meanAREAPositiveCells, stdDevAREAPositiveCells);

if(NegCell!=0){

//Create array with only data from negative cells detected

CELLNegativeMeanData=newArray(NegCell);

CELLNegativeAreaData=newArray(NegCell);

CELLNegativeROIName=newArray(NegCell);

positionarray=0;

for(data=0;data<myCellsROIMean.length;data++){

if(CELLPositiveORNegativeSignal[data]=="Negative"){

CELLNegativeMeanData[positionarray]=myCellsROIMean[data];

CELLNegativeAreaData[positionarray]=myCellsROIArea[data];

CELLNegativeROIName[positionarray]=myCellsROINames[data];

positionarray++;

}

}

//Create only negative cells data Results table

Array.show("Negative_Cells_Data_Results", CELLNegativeROIName, CELLNegativeAreaData,CELLNegativeMeanData);

selectWindow("Negative_Cells_Data_Results");

saveAs("Results", results+myimagename+"_Ch_"+AnalysisChannelLabel+"_Negative_Cells_Data_Results.xls");

selectWindow(myimagename+"_Ch_"+AnalysisChannelLabel+"_Negative_Cells_Data_Results.xls");

run("Close");

//Get statistics of only negative cells array

Array.getStatistics(CELLNegativeMeanData, dataIdonotneed, dataIdonotneed, meanMEANNegativeCells, stdDevMEANNegativeCells);

Array.getStatistics(CELLNegativeAreaData, dataIdonotneed, dataIdonotneed, meanAREANegativeCells, stdDevAREANegativeCells);

//Save final Cell ROIs

roiManager("deselect");

roiManager("save", results+myimagename+"-Cells_ROIs.zip");

roiManager("reset");

}else{

print("No positive cells detected");

meanPositiveCells=0;

stdDevPositiveCells=0;

}

close("CELLP");

close(CellChannelLabel);

}

}

if (NucleiAnalysis=="yeah"){

//****************Begining of NUCLEI analysis*************************************************************

//Nuclei channel image pre-processing

selectImage(ORI);

run("Select None");

run("Duplicate...", "title="+NucChannelLabel+" duplicate channels="+Nucch);

NUCS=getImageID();

selectImage(NUCS);

run("Gaussian Blur...", "sigma=3");

//Nuclei Detection using StarDist plugin

run("Command From Macro", "command=[de.csbdresden.stardist.StarDist2D], args=['input':'"+NucChannelLabel+"', 'modelChoice':'Versatile (fluorescent nuclei)', 'normalizeInput':'true', 'percentileBottom':'25.700000000000003', 'percentileTop':'99.8', 'probThresh':'0.4', 'nmsThresh':'0.1', 'outputType':'ROI Manager', 'nTiles':'10', 'excludeBoundary':'2', 'roiPosition':'Automatic', 'verbose':'false', 'showCsbdeepProgress':'false', 'showProbAndDist':'false'], process=[false]");

//Rename final ROIs	using macro function

var allmyNUCrois=roiManager("count");

myNUCsROINames=RENAMEmyROIs(0, allmyNUCrois, "N", "white");

roiManager("deselect");

//Measure NUC ROI parameters on channel of interest

selectImage(ANALYSIS);

run("Select None");

myNUCsROIArea=MEASUREmyROIs(0, allmyNUCrois, "Area", 0);

myNUCsROIMean=MEASUREmyROIs(0, allmyNUCrois, "Mean", 0);

//Create Nuclei total data Results table

Array.show("Nuclei_Data_Results", myNUCsROINames, myNUCsROIArea, myNUCsROIMean);

selectWindow("Nuclei_Data_Results");

saveAs("Results", results+myimagename+"_Ch_"+AnalysisChannelLabel+"_Nuclei_Data_Results.xls");

selectWindow(myimagename+"_Ch_"+AnalysisChannelLabel+"_Nuclei_Data_Results.xls");

run("Close");

//Obtain threshold for positive signal Nuclei classification (% of Max Signal found will be used)

////Get array statistics to find max value

Array.getStatistics(myNUCsROIMean, minMeanNUCValue, maxMeanNUCValue, meanMeanNUCValue);

print("Max mean value of all nuclei detected "+ maxMeanNUCValue);

print("Mean mean value of all nuclei detected "+ meanMeanNUCValue);

print("Min mean value of all nuclei detected "+ minMeanNUCValue);

ThresholdValueNUCs=(maxMeanNUCValue*(PercentageMaxSignalNucCh/100));

DifferenceOfSignalsNucs=meanMeanNUCValue*PercentageDifferenceSignalNuclei;

print("Threshold value used for positive nuclei classification on channel "+ AnalysisChannelLabel+" will be >= to "+ ThresholdValueNUCs);

print("Second Threshold value used for positive cell classification on channel "+ AnalysisChannelLabel+" will be > to "+ DifferenceOfSignalsNucs);

//Nuclei classification

////New array for positive/negative labeling of Nuclei. Set counters

NUCPositiveORNegativeSignal=newArray(allmyNUCrois);

PosNUC=0;

NegNUC=0;

for(data=0;data<myNUCsROIMean.length;data++){

if(myNUCsROIMean[data]>=ThresholdValueNUCs && myNUCsROIMean[data]>DifferenceOfSignalsNucs){

NUCPositiveORNegativeSignal[data]="Positive";

PosNUC++;

roiManager("select", data);

roiManager("rename", "+Nuc_"+data+1);

roiManager("Set Color", "Magenta");

roiManager("Set Line Width", 2);

}else{

NUCPositiveORNegativeSignal[data]="Negative";

NegNUC++;

roiManager("select", data);

roiManager("rename", "-Nuc_"+data+1);

roiManager("Set Color", "Cyan");

roiManager("Set Line Width", 2);

}

}

if(PosNUC!=0){

//Create array with only data from positive nuclei detected

NUCPositiveMeanData=newArray(PosNUC);

NUCPositiveAreaData=newArray(PosNUC);

NUCPositiveROIName=newArray(PosNUC);

positionarray=0;

for(data=0;data<myNUCsROIMean.length;data++){

if(NUCPositiveORNegativeSignal[data]=="Positive"){

NUCPositiveMeanData[positionarray]=myNUCsROIMean[data];

NUCPositiveAreaData[positionarray]=myNUCsROIArea[data];

NUCPositiveROIName[positionarray]=myNUCsROINames[data];

positionarray++;

}

}

//Create only positive nuclei data Results table

Array.show("Positive_Nuclei_Data_Results", NUCPositiveROIName, NUCPositiveAreaData,NUCPositiveMeanData);

selectWindow("Positive_Nuclei_Data_Results");

saveAs("Results", results+myimagename+"_Ch_"+AnalysisChannelLabel+"_Positive_Nuclei_Data_Results.xls");

selectWindow(myimagename+"_Ch_"+AnalysisChannelLabel+"_Positive_Nuclei_Data_Results.xls");

run("Close");

//Get statistics of only positive nuclei array

Array.getStatistics(NUCPositiveMeanData, dataIdonotneed, dataIdonotneed, meanMEANPositiveNUCs, stdDevMEANPositiveNUCs);

Array.getStatistics(NUCPositiveAreaData, dataIdonotneed, dataIdonotneed, meanAREAPositiveNUCs, stdDevAREAPositiveNUCs);

}else{

print("No positive nuclei found");

meanMEANPositiveNUCs=0;

stdDevMEANPositiveNUCs=0;

meanAREAPositiveNUCs=0;

stdDevAREAPositiveNUCs=0;

}

if(NegNUC!=0){

//Create array with only data from Negative nuclei detected

NUCNegativeMeanData=newArray(NegNUC);

NUCNegativeAreaData=newArray(NegNUC);

NUCNegativeROIName=newArray(NegNUC);

positionarray=0;

for(data=0;data<myNUCsROIMean.length;data++){

if(NUCPositiveORNegativeSignal[data]=="Negative"){

NUCNegativeMeanData[positionarray]=myNUCsROIMean[data];

NUCNegativeAreaData[positionarray]=myNUCsROIArea[data];

NUCNegativeROIName[positionarray]=myNUCsROINames[data];

positionarray++;

}

}

//Create only Negative nuclei data Results table

Array.show("Negative_Nuclei_Data_Results", NUCNegativeROIName, NUCNegativeAreaData,NUCNegativeMeanData);

selectWindow("Negative_Nuclei_Data_Results");

saveAs("Results", results+myimagename+"_Ch_"+AnalysisChannelLabel+"_Negative_Nuclei_Data_Results.xls");

selectWindow(myimagename+"_Ch_"+AnalysisChannelLabel+"_Negative_Nuclei_Data_Results.xls");

run("Close");

//Get statistics of only Negative nuclei array

Array.getStatistics(NUCNegativeMeanData, dataIdonotneed, dataIdonotneed, meanMEANNegativeNUCs, stdDevMEANNegativeNUCs);

Array.getStatistics(NUCNegativeAreaData, dataIdonotneed, dataIdonotneed, meanAREANegativeNUCs, stdDevAREANegativeNUCs);

//Save final Nuclei ROIs

roiManager("deselect");

roiManager("save", results+myimagename+"-Nuclei_ROIs.zip");

roiManager("reset");

}else{

print("No Negative nuclei found");

meanMEANNegativeNUCs=0;

stdDevMEANNegativeNUCs=0;

meanAREANegativeNUCs=0;

stdDevAREANegativeNUCs=0;

}

}

//Create Summary Results table (cells and nuclei)

run("Clear Results");

if (isOpen("TempResults")){Table.rename("TempResults", "Results");}

setResult("Image", myposition, myimagename);

setResult("Channel for analysis", myposition, "# Ch:"+Analysisch+", "+AnalysisChannelLabel);

setResult("Area units ^2", myposition, unitvoxel);

if (CellAnalysis=="yeah"){

setResult("# Total Cells detected", myposition, allmyCELLrois);

setResult("# +Cells detected", myposition, PosCell);

setResult("# -Cells detected", myposition, NegCell);

setResult("Ratio +Cells/Total detected", myposition, (PosCell/allmyCELLrois));

setResult("Mean of IntensityMean of +Cells", myposition, meanMEANPositiveCells);

setResult("STD of IntensityMean of +Cells", myposition, stdDevMEANPositiveCells);

setResult("Mean of Area of +Cells", myposition, meanAREAPositiveCells);

setResult("STD of Area of +Cells", myposition, stdDevAREAPositiveCells);

setResult("Mean of IntensityMean of -Cells", myposition, meanMEANNegativeCells);

setResult("STD of IntensityMean of -Cells", myposition, stdDevMEANNegativeCells);

setResult("Mean of Area of -Cells", myposition, meanAREANegativeCells);

setResult("STD of Area of -Cells", myposition, stdDevAREANegativeCells);

}

setResult("----", myposition, "	");

if (NucleiAnalysis=="yeah"){

setResult("# Total Nuclei detected", myposition, allmyNUCrois);

setResult("# +Nuclei detected", myposition, PosNUC);

setResult("# -Nuclei detected", myposition, NegNUC);

setResult("Ratio +Nuclei/Total detected", myposition, (PosNUC/allmyNUCrois));

setResult("Mean of IntensityMean of +Nuclei", myposition, meanMEANPositiveNUCs);

setResult("STD of IntensityMean of +Nuclei", myposition, stdDevMEANPositiveNUCs);

setResult("Mean of Area of +Nuclei", myposition, meanAREAPositiveNUCs);

setResult("STD of Area of +Nuclei", myposition, stdDevAREAPositiveNUCs);

setResult("Mean of IntensityMean of -Nuclei", myposition, meanMEANNegativeNUCs);

setResult("STD of IntensityMean of -Nuclei", myposition, stdDevMEANNegativeNUCs);

setResult("Mean of Area of -Nuclei", myposition, meanAREANegativeNUCs);

setResult("STD of Area of -Nuclei", myposition, stdDevAREANegativeNUCs);

}

updateResults();

selectWindow("Results");

saveAs("Results", results+"MyResults.xls");

saveAs("Text", results+"MyResults.txt");

Table.rename("Results","TempResults");

myposition++;

run("Close All");

//Save Log

if(isOpen("Log")){

selectWindow("Log");

saveAs("Text", results+"Log.txt");

}

}

}

}

waitForUser("Analysis done, your results are here:"+results);

}

//------------------------

//MACRO FUNCTIONS:

//------------------------

//1)

////*****Function to rename and change colour of rois in roi manager****

//Output of the function is an array containing roi names.

//Variables to feed the function:

//*Begin,end (range of rois to be taken into account for the function)

//*roiname (name that will be applied to rois);

//*roicolor(color applied to rois)

function RENAMEmyROIs(begin, end, roiname, roicolor){

myarrayelements=end-begin;

mynamesarray=newArray(myarrayelements);

position=0;

for(n=begin; n<end;n++){

roiManager("select", n);

roiManager("rename", roiname+"-"+position+1);

roiManager("Set Color", roicolor);

roiManager("Set Line Width", 1);

roiManager("select", n);

mynamesarray[position]=Roi.getName;

position++;

}

roiManager("deselect");

return mynamesarray;

}

//2)

//****Function to measure any desired parameter in a range of rois from roi manager (using or not the limit to threshold option)****

//Output of the function is an array containing parameter values for all rois in range

//Variables to feed the function:

//*begin,end (range of rois to be taken into account)

//*parameter (parameter to measure, any parameter that can be obtained from the Set Measurements list)

//*thresholdYESorNO (use 1 to limit measurements to a threshold, use 0 to measure parameter without threshold)

function MEASUREmyROIs(begin, end, parameter, thresholdYESorNO){

myarrayelements=end-begin;

mymeasurementsarray=newArray(myarrayelements);

position=0;

for(n=begin; n<end;n++){

roiManager("select", n);

if(thresholdYESorNO==1){

List.setMeasurements("limit");

}else{ List.setMeasurements;}

mymeasurementsarray[position]=List.getValue(parameter);

position++;

}

roiManager("deselect");

return mymeasurementsarray;

}
